# Risk to European birds from collisions with wind-energy facilities

**DOI:** 10.1101/2025.11.24.685024

**Authors:** Adrienne Etard, Martin Jung, Piero Visconti

## Abstract

In line with Europe’s decarbonization goals, wind-power capacity is projected to increase in future years. However, wind-energy facilities can affect biodiversity; flying animals can fatally collide with wind-energy infrastructure. Here, we assessed the reported risk posed by collisions to 108 European birds, comparatively across species. We drew risk maps to assess current hotspots of risk. We employed a customised risk approach, considering that risk emerged from the interaction between (1) reported impacts (the total estimated number of fatalities at the species-level, accounting for current exposure to wind turbines) and (2) vulnerability to these impacts (the degree to which species may be affected by collision mortality). We used a quantitative synthesis of fatality numbers at wind-energy facilities to quantify collision-mortality rates at the species-level. We derived (1) by further combining these with information on species suitable areas and on current wind-turbine deployment. We estimated species’ vulnerability from ecological characteristics (generation length, clutch size, and estimates of European suitable area) assumed to reflect species’ ability to cope with disturbances. Overlapping vulnerability with estimated impacts, we classified species into different risk categories, considering species to be at higher relative risk when more vulnerable and more impacted. We assessed where species falling into the risk categories might occur, and where possible conflicts with wind-energy deployment might arise. We found several risk hotspots notably located in the Iberian Peninsula and Northern Europe. Our work helps inform wind-power deployment and spatial planning at the European scale with the aim of minimising negative biodiversity impacts.

## 1. Introduction

With the 2021 adoption of the European Climate Law, the European Union has committed to becoming climate-neutral by 2050, i.e., to reaching net-zero greenhouse gas emissions. Key to the EU’s decarbonisation is the transformation of energy systems, most notably the reduction of fossil fuels and increases in the share of renewable energy sources. In 2022, wind power alone accounted for about 16% of the electricity demand in the EU (Wind Europe, 2022). In line with the European Climate Law, the installed capacity needs to increase by more than 100% to meet the 2030 targets (to reach a total capacity of 440 GW by 2030 with regards to the REPower EU energy target; Wind Europe, 2022).

While wind energy is a key sector for renewable energy production, wind-energy facilities can have adverse impacts on biodiversity (Loss et al., 2013; Schöll & Nopp-Mayr, 2021). These impacts can be an important barrier to the social acceptance of wind power (Voigt et al., 2019; Vuichard et al., 2022). Negative biodiversity impacts can be attributed to land-use change and habitat disturbance occurring from the construction phase of the facilities and necessary structures (e.g., roads and power lines). Habitat disturbance can lead to the direct displacement of species through the loss of suitable habitat (Marques et al., 2020), and to reductions in local abundance and population densities (Fernández-Bellon et al., 2019). For flying animals, wind turbines may present a barrier inducing avoidance behaviour and altering migratory routes (Cabrera-Cruz & Villegas-Patraca, 2016; Santos et al., 2022a). Wind-energy facilities also constitute a collision hazard for flying animals, birds and bats in particular (Smallwood, 2013; Thaxter et al., 2017), but also insects (Voigt, 2021). Collision mortality can be of concern for the long-term viability of animal populations (Duriez et al., 2023; Gómez-Catasús et al., 2018; May et al., 2019). Of particular concern are species that cumulate multiple factors rendering them vulnerable to increased mortality, such as low population densities, slow pace of life, and/or species that are already threatened (Carrete et al., 2009; Desholm, 2009; Kuvlesky Jr. et al., 2010). For instance, large soaring birds have been a group of interest since their reliance on wind resources to gain altitude (thermal and orographic updrafts in particular) can conflict with areas favourable for wind-energy developments (Farfán et al., 2023; Santos et al., 2022b; Smeraldo et al., 2020) while poor flight manoeuvrability can make them prone to colliding, and low reproductive outputs can make the species more vulnerable to added mortality (Bellebaum et al., 2013; Dahl et al., 2013). Assessing the risk posed by collision mortality to different species is therefore important to inform spatial deployment and minimise biodiversity impacts.

Past research has shown that collision mortality is influenced by many factors, which can be broadly classified into three categories (Marques et al., 2014): (1) site-specific factors that relate to local landscape configuration and environmental features, such as wind speed, weather, landforms, habitat availability for different species, etc; (2) characteristics of the wind-energy facilities, such as spatial configuration, turbine capacity, lighting, etc; and (3) species characteristics underlying vulnerability (e.g. seasonal occurrence, morphology, behaviour, etc. (May et al., 2015)). Collision mortality is influenced by complex interactions among these factors, which points to an important degree of context-specificity (Drewitt & Langston, 2008; Marques et al., 2014). Consequently, the siting of wind-energy facilities and their spatial configuration are important to reduce collision mortality (Carrete et al., 2012; Schaub, 2012). However, in a quantitative synthesis of collision-mortality data, Thaxter et al. (2017) notably showed that species’ habitat preferences and dispersal distances were significantly associated with collision mortality. Using these associations, Thaxter et al. (2017) estimated the vulnerability of birds to collision at global scales, inferring collision-mortality rates for species for which there were no collision-fatality records. However, they did not investigate whether landscape characteristics were associated with collision risk; and did not consider the degree of exposure of different species to wind power.

Here, building upon Thaxter et al. (2017), we aimed to assess the risk to European birds from collision mortality, and to provide a spatial assessment of risk for Europe. Our main objective was to derive risk maps at the European scale that could be integrated into decision-making tools and inform spatial deployment. To this end, we drew from past risk and vulnerability approaches (Bellard et al., 2024; Foden et al., 2013; Intergovernmental Panel on Climate Change (2007, 2014)). We devised a custom risk approach considering that risk emerged from the interaction between species intrinsic vulnerability and from collision-mortality impacts (estimated total number of fatalities), which accounted for current exposure to wind turbines (Fig. 1). We estimated these two dimensions of risk (impacts and vulnerability) separately, thus independently deriving an ‘impact index’ and a ‘vulnerability index’ at the species level.

**Fig. 1.**
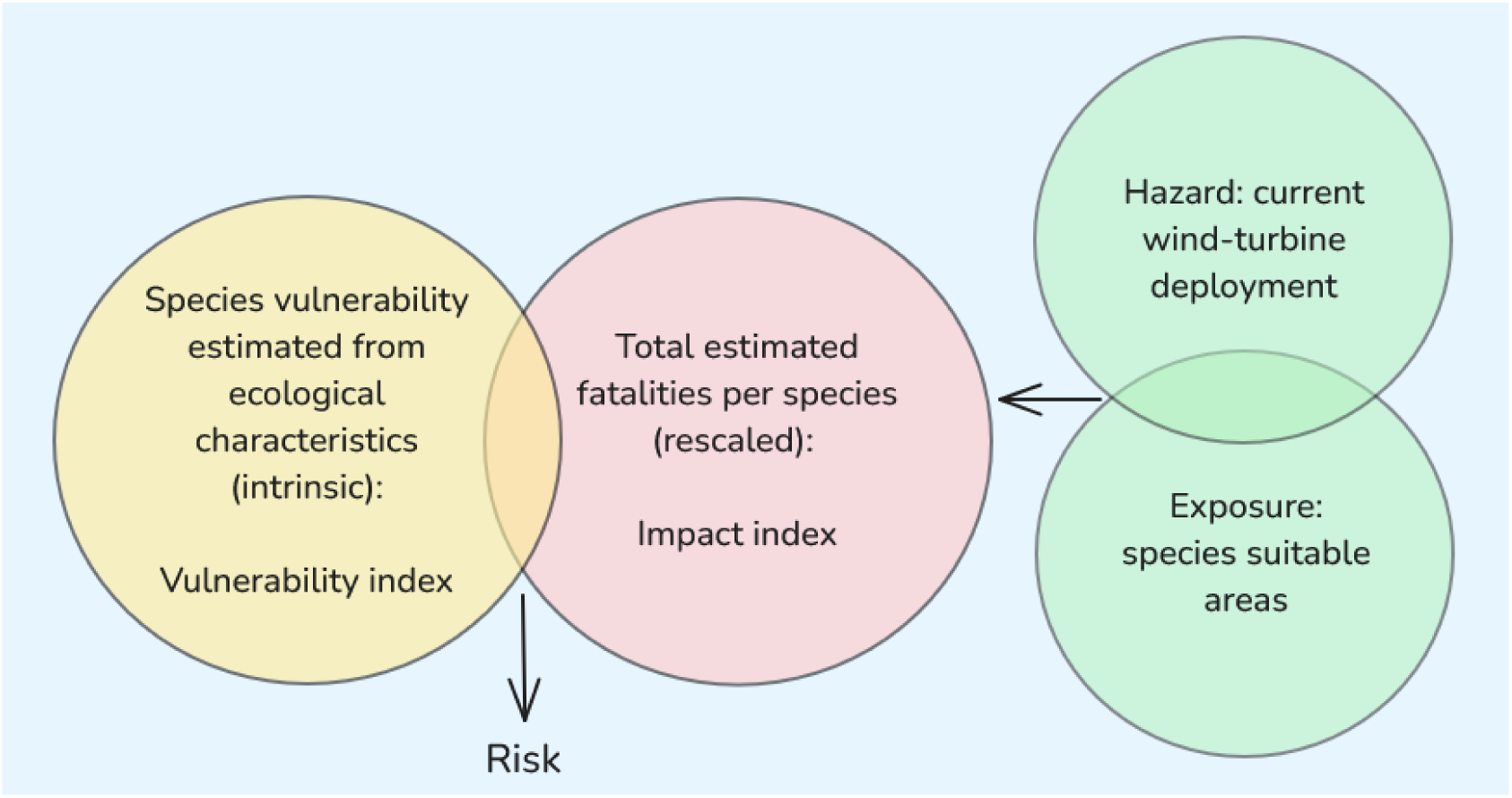
Conceptual and terminological clarifications on risk and vulnerability. Foden et al. (2019) described a change in the terminology relating to vulnerability and risk between the 4^th^ and the 5^th^ Intergovernmental Panel on Climate Change (IPCC) reports. In the IPCC 4^th^ assessment (Intergovernmental Panel on Climate Change, 2007), vulnerability emerges from the interaction of sensitivity (degree to which a system is likely to be affected, intrinsic), adaptive capacity (ability of a system to adjust, intrinsic), and exposure (degree of disturbance experienced by a system, extrinsic). Vulnerability defined as such is widely employed in the field of conservation biology for vulnerability assessments (e.g. Bellard et al., 2024; Foden et al., 2013). The IPCC 5^th^ assessment report (Intergovernmental Panel on Climate Change, 2014) presented an alternative risk framework, with differences in terminology. Risk results from the interaction of vulnerability (considered intrinsic: propensity of a system to be negatively affected; encompasses notions of sensitivity and adaptive capacity), exposure (considered intrinsic: presence of system of interest in places that could be affected), and hazard (extrinsic: presence of a particular physical threat). For further clarifications, see Foden et al. (2019). Here, we built upon these frameworks to customize a risk approach. We considered risk as emerging from the interaction between vulnerability (intrinsic ability of species to cope with disturbances, here degree to which species might be able to withstand the additive mortality) and impacts, which we defined as the total estimated number of collision fatalities at the species level, accounting for the current exposure of species to wind turbines. Note that our definition of ‘impact’ therefore differs from that of the IPCC 5^th^ assessment report (Foden et al., 2019).

First, we estimated the impact index using the data from Thaxter et al. (2017). In doing so, another important question arose: whether we could consider species for which there were no fatality records in our analyses, by estimating collision-mortality rates for such species from associations with species-level characteristics, following Thaxter et al. (2017). Therefore, a secondary objective of our study was to investigate whether collision-mortality rates were associated with species- and site-level characteristics, building upon Thaxter et al. (2017). Our rationale was that, should collision mortality be associated with these characteristics, we could estimate it based on such associations (Thaxter et al. 2017), therefore expanding the risk estimation to all European species, including those that were not in the data (rather than focussing only on species for which there were reported fatalities). Thus, within our risk framework, we investigated whether species- and site-level characteristics were associated with collision-mortality rates using statistical models, and whether such characteristics outperformed taxonomy only in explaining the variation in collision-mortality rates. We further used the outputs of the models to estimate the impact index, combining estimated collision-mortality rates with information on wind-turbine deployment in Europe and on species’ likely habitat suitability (i.e., exposure; Fig. 1).

Second, we developed a vulnerability index that accounted for additional ecological characteristics reflecting species’ likely intrinsic vulnerability to disturbances. Finally, we intersected the impact index with the vulnerability index to assess the risk across species. In line with vulnerability assessments, our approach was therefore comparative, reflecting the risk of the considered species relative to one another. As wind power is likely to expand in Europe in future years, and as species are unevenly affected by wind-energy facilities, assessing which factors influence collision mortality and evaluating which species and regions might be at more risk can help putting into place appropriate mitigation measures.

## 2. Materials and methods

### 2.1. Overview of the workflow

Fig. S1 describes the conceptual framework and objectives for this work. Our aim was to assess the relative risk for European species, based on an estimation of vulnerability and of estimated impacts. A key consideration was whether we could extend the risk estimation to species for which there were no fatality records. Thus, a secondary aim was to assess whether collision-mortality rates were associated with species- and site-level characteristics, therefore investigating if collision-mortality rates could be estimated for all European species based on such possible associations (decision tree on Fig. S1; Section 2.2). The methodology behind the risk assessment (Section 2.3) was therefore partly informed by the Results from 2.2. in defining the taxonomic scope and modelling approach. In brief, we (1) estimated an index of collision-mortality impacts across species; impacts were quantified as the total number of fatalities estimated for each species. This index accounted for exposure and hazard (wind turbines overlapping with species suitable habitats), combined with estimated collision-mortality rates at the species level to estimate fatality numbers (2.2). (2) We built the vulnerability index by considering species’ life history (generation length and clutch size) and rarity (European suitable range area), assuming such characteristics to be proxies for vulnerability, expressed as the species’ ability to withstand the additional collision mortality. The rationale was that species with longer generation lengths and/or with fewer offsprings, and (geographically) rarer species, might be more vulnerable because, from a demographic point of view, it might take such species longer to recover from decreases in population size than species with shorter generation lengths, a larger clutch size, or than more (geographically) common species (Foden et al., 2019). (3) Finally, combining the impact index with the vulnerability index (Hyman et al., 2025), we assessed the comparative, relative risk across species and assessed which European regions harboured species at higher risk.

### 2.2. Are species- and site-level characteristics associated with collision-mortality rates?

#### 2.2.1. Fatality-count data

We used data from Thaxter et al. (2017), in which peer-reviewed and non-peer-reviewed studies and reports that estimated collision mortality at wind-energy facilities were compiled. Thaxter et al. (2017) identified 88 sources for birds suitable for inclusion, spanning locations biased towards Europe and North America. The studies were heterogenous, e.g. varying in the number of wind turbines, sampling year(s), or search protocols. Thaxter et al. (2017) compiled estimates at the wind-energy facility level, also reporting turbine capacity, number of wind turbines, central location of the facility (central longitude and latitude of turbines), and study duration. In addition, Thaxter et al. (2017) estimated the quality of each study, e.g., whether studies corrected for carcass detectability (Huso & Dalthorp, 2014; Nilsson et al., 2023; Domínguez del Valle et al., 2020), characterising study quality as “very low”, “low”, “medium”, or “high”, depending on the application of correction factors. For further methodological details, see Thaxter et al. (2017).

Over 90% of the records reported either (annual) collision-mortality rates or fatality counts, depending on the study. We subset the data to include only such records (e.g., described as “annual mortality per turbine”, “total fatalities”) and for which study duration and number of turbines were known. We standardised the values across studies to obtain annual mortality rates throughout (i.e., number of fatalities per year-turbine). These rates were converted back to count data prior to fitting the models, as required by Poisson models (thus we refer to these data to as ‘fatality-count data’; Supporting Information S2; Fig. S2).

Although our assessment focussed on European species, we considered sites located in other regions (e.g. North America), because we had no expectation that ecological characteristics should influence collision mortality differently in different regions. In addition, some species were sampled across several continents, so retaining all sites increased sample size. Despite this, the data remained biased, not only geographically (Fig. S3), but also taxonomically, with some species sampled only a few times (Fig. S4; 66% of the species were recorded at least twice, 47% at least three times, and 16% more than 10 times).

#### 2.2.2. Investigating associations with species- and site-level characteristics

We combined the fatality-count data with species-level ecological characteristics (Table 1a), targeting traits that related to movement (i.e., hand-wing index, migratory status) and species habitat preferences (Thaxter et al. 2017).

**Table 1.**
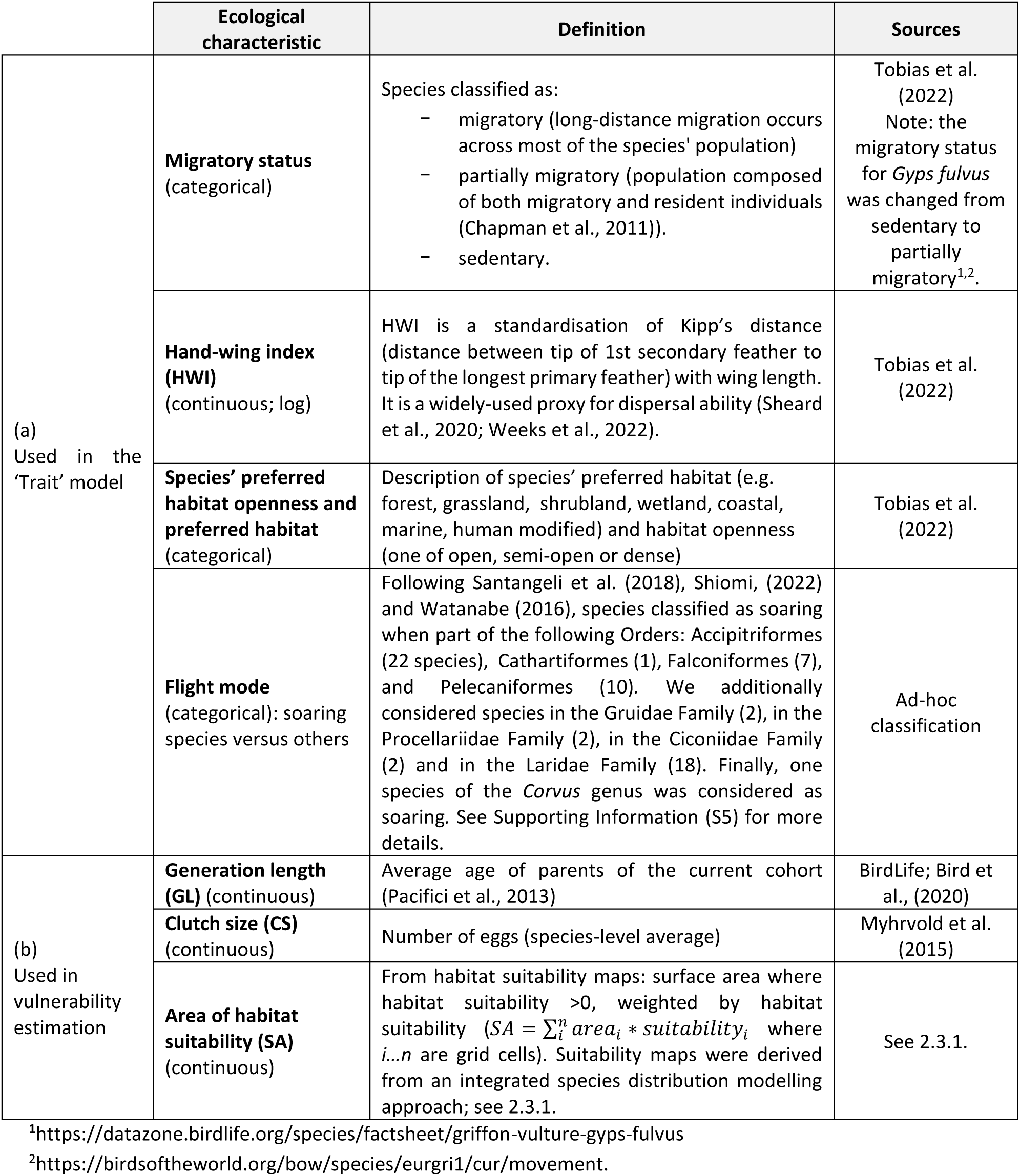
Ecological characteristics considered (a) in the models investigating associations between species-level characteristics, site-level characteristics, and collision-mortality rates; (b) in the comparative assessment of species vulnerability: definitions and sources.

At the central location of the wind-energy facilities, we obtained four environmental characteristics: mean wind speed at 100 m, elevation, terrain roughness, and topographic index. Roughness reflected the variability in elevation around a location, while topographic index reflected whether a location was relatively lower or higher than its surroundings. We obtained mean wind speed from the Global Wind Atlas (https://globalwindatlas.info/en; data downloaded 23/10/2023 and aggregated at 1 x 1 km using an arithmetic mean). Elevation, roughness, and topographic index were derived from a global digital elevation model (Amatulli et al. (2018); http://www.earthenv.org/topography). We used different buffer sizes (0, 10, 100, 500, and 1000 m) to capture the landscape context around each location (extracting values around the locations falling within each buffer area, then using the mean across the buffer cells), but since the estimations were overall congruent across buffer sizes (Fig. S5) we used a buffer of 1 km around each location throughout.

First, we investigated whether species’ ecological characteristics (Table 1a), site-level characteristics, turbine capacity and study quality were associated with collision-mortality rates. Study quality was specifically included to control for the degree to which corrections for carcass detectability were applied across studies (Thaxter et al., 2017). We fitted generalized linear mixed-effects models, using a Bayesian framework implemented with the R-package ‘brms’ (Bürkner, 2017, 2018, 2021). We fitted fatality counts (per record within each study) as the response, using a zero-inflated Poisson error distribution to account for excess-zeroes, using an offset variable to control for study duration and varying number of turbines within wind-energy facilities. Random intercepts accounted for variation in experimental design across studies and for the spatial structuring of sites and studies; another random intercept accounted for the identity of species, controlling for repeated observations and taxonomic non-independence. We fitted an observation-level random effect to control for overdispersion (Harrison, 2014). Drawing on Thaxter et al. (2017)’s findings that species with intermediate dispersal distances had significantly higher collision-mortality rates, and on the association between hand-wing index and dispersal (Sheard et al., 2020; Weeks et al., 2022), we treated hand-wing index as a continuous predictor with a quadratic effect. In the first iteration of this model (referred to as ‘trait model’), we considered the effects of habitat openness, but we further tested if using habitat categories instead of openness (Table 1a) improved the model. The initial trait model considered the effects of mean wind speed, and we further tested whether adding the other site-level variables (terrain roughness, elevation, topographic index) improved the model.

Second, we fitted an alternative model, whereby species identity was fitted as a fixed effect, while other species- and site-level characteristics were removed (except for turbine capacity and study quality). This model (referred to as ‘taxonomic model’) was used to assess whether species- and site-level characteristics outperformed taxonomy in explaining the variation in collision-mortality rates across species.

Each model was fitted through Markov-Chain Monte Carlo, using 4 chains, 5000 warm-up iterations, 50,000 sampling iterations, and thinning intervals of 10 with weakly informative priors to improve sampling convergence. We compared all six models using the ‘*loo_compare’* function of the brms package, using ‘elpd’ differences (Vehtari et al., 2017).

### 2.3. Risk estimation

#### 2.3.1. Impact index

We used the database from Fischereit et al. (in preparation), in which publicly available and commercial datasets (including https://www.thewindpower.net/) were combined to create a European wind-turbine database. We combined wind-turbine locations and capacity (i.e., hazard; Fig. 1) with species habitat suitability maps (i.e., exposure; Fig. 1) and with the fitted models to quantify the total number of fatalities for each species within their suitable habitat (i.e., impact index). For each species, a habitat suitability map was estimated using the modelling approach developed by Jung (2023). Suitability maps were derived from an integrated species distribution modelling approach, combining a range of biodiversity datasets (from GBIF, eBIRD and Natura 2000) and environmental variables (e.g., climate and land-cover variables) into an ensemble-modelling framework to predict how suitable the local environmental conditions might be for a species. A weighted ensemble prediction from multiple ‘engines’ was calculated and a threshold based on the True Skill Statistics obtained. For each ensemble prediction, values below the threshold were set to 0, and we considered as suitable all pixels where suitability was above the threshold. More details about the maps can be found in the supplementary materials of Chapman et al. (2023) and in Visconti et al. (2024) in which similar maps were used. All available maps were projected with the Lambert azimuthal equal-area projection, with a resolution of 10 km by 10 km.

In each suitable grid-cell where turbines were detected, fatality counts were estimated as a function of species identity and turbine capacities, using the taxonomic model (which had a similar predictive performance to the trait model; see Results), and therefore not expanding the analysis to species for which there were no fatality records (Fig. S1). Estimations were obtained for the highest study quality and were made marginal to location (site) and study identity. To this end, we linearised the relationship between turbine capacity and fatality counts for each species, instead of using the exponential relationship estimated from the Poisson model. We did this because the turbine capacity in the fitted data ranged between 0.015MW and 2.5MW, while turbines with a capacity of up to 8MW occurred in Europe. Assuming an exponential relationship would lead to unrealistic estimations of fatality counts for large capacities (Fig. S6). For each species, we used the model to predict values for capacities of 0.015MW and 2.5MW (range of values in the fitted data) and estimated the slope assuming a linear relationship. We then used the estimated slopes to predict fatality counts given turbine capacities, weighting the estimation by the suitability value of each relevant grid cell. We thus derived estimations of fatality counts per year across all turbines falling within the suitable habitat of each species (i.e., total fatalities per year). The impact index was then rescaled between 0 and 1 across species, to provide a relative impact index with higher values reflecting higher relative impacts and lower values reflecting lower relative impacts.

#### 2.3.2. Vulnerability index

The vulnerability index accounted for three ecological characteristics: generation length (GL), clutch size (CS), and area of habitat suitability (SA; Table 1b). Drawing from vulnerability analyses (Bellard et al., 2024), the vulnerability index was calculated as a composite index, by summing of the characteristic values, after rescaling each characteristic across species for comparability, as GL_rescaled_ + (1-CS_rescaled_) + (1-SA_rescaled_). We took the opposite of CS and SA to reflect the higher vulnerability of species with smaller clutch sizes or areas of habitat suitability, compared to species with larger clutch sizes or areas of habitat suitability. The vulnerability dimensions were not too highly correlated (Table S1). We then rescaled the final vulnerability index, so that it ranged between 0 and 1 with lower values reflecting lower relative vulnerability and higher values reflecting higher relative vulnerability.

#### 2.3.3. Risk categories

We assessed species risk by considering the relative position of species in the risk space (vulnerability index against impact index; Fig. 2). We considered species to be at ‘high risk’ when exhibiting higher relative vulnerability and higher relative impacts, species to be at ‘high latent risk’ when exhibiting higher relative vulnerability but lower relative impacts (meaning that species could be negatively affected if more impacted), species to be ‘possible persisters’ when exhibiting lower relative vulnerability with higher relative mortality impacts (species possibly able to persist despite high impacts), and species to be ‘possible adapters’ when exhibiting lower relative vulnerability with lower relative mortality impacts (species that currently have low mortality impacts, but could be able persist if cumulative collision mortality increased; note that the terms were adapted from Foden et al. (2013) for the purpose of this work and of defining risk categories here). In absence of specific demographic information to set ecologically-informed thresholds, we used the 75% quantiles of the distributions of the impact index and of the vulnerability index as threshold values, considering species to be highly vulnerable and highly impacted when falling above these quantiles (overlapping top 25% species in both dimensions). Because using different threshold values could influence the results, we also considered 50% quantiles to investigate the congruence of the results. We thus classified species into the different risk categories (Fig. 2).

**Fig. 2.**
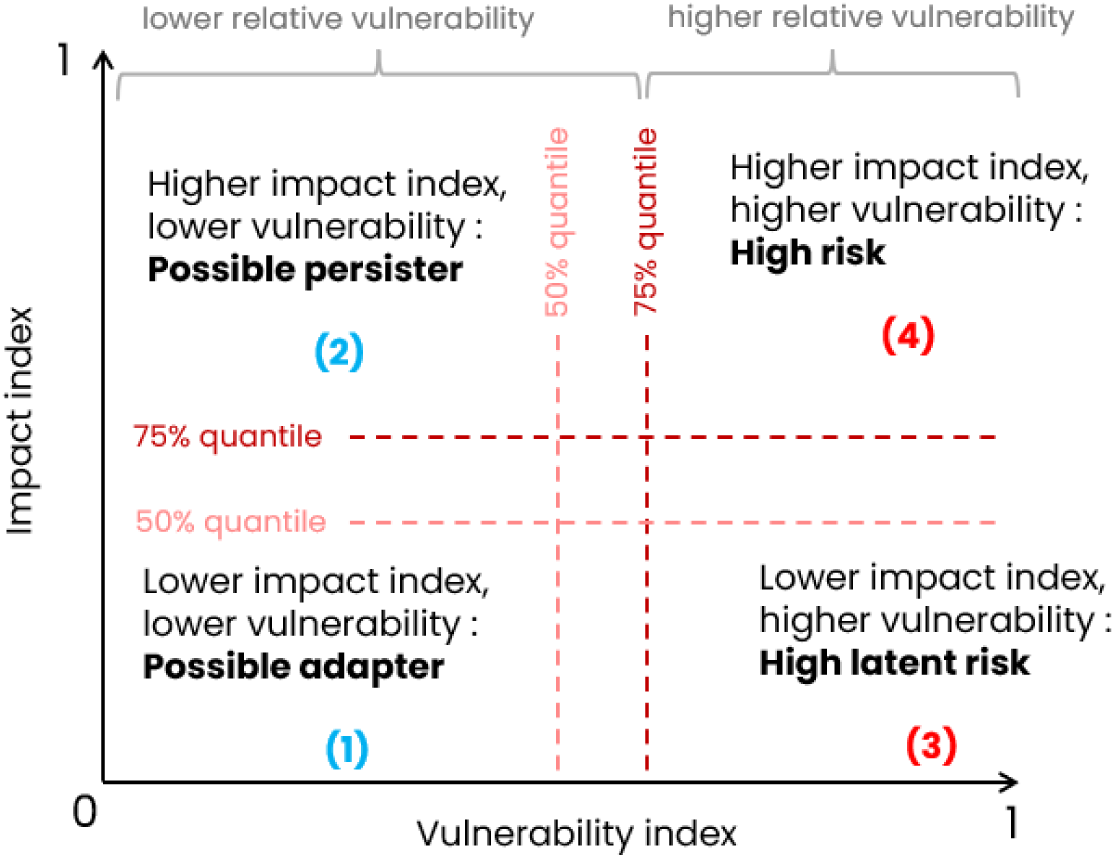
We derived a risk category at the species-level by considering the overlap between the vulnerability index and the impact index. Terminology adapted from Foden et al. (2013). The 75% quantile of each distribution was used as a threshold above which we considered species to be highly vulnerable and highly impacted (species classified as “high risk” when falling above both thresholds), and we used 50% quantiles to check the congruence of the results when using lower threshold values. We also highlight relative vulnerability groupings (i.e. grouping species into lower/higher relative vulnerability based on the chosen quantile).

#### 2.3.4. Mapping risk

To assess hotspots of risk, we mapped pseudo-species richness in each risk category, for the two threshold values applied to classify species into the risk categories (75% and 50%). We also grouped high risk and high latent risk species on the one hand; and possible persisters and possible adapters on the other hand (i.e., lower versus higher vulnerability groupings). Pseudo-richness maps were derived by stacking the suitability maps.

Further, combining the richness maps with the current wind-turbine locations, we assessed the spatial overlap between wind-turbine locations and local pseudo-richness in the different risk categories and groupings. We derived bivariate plots highlighting areas with both high turbine densities and a relatively higher number of species in each group (‘biscale’ package, Prener et al. (2022)). Turbine kernel densities were derived using the ‘*st_kde*’ function of the ibis.iSDM package (Jung, 2023).

All data processing and analyses were conducted with R v.4.2.3 (R Core Team). The output pseudo-species richness maps were made available at 10.5281/zenodo.14717411.

## 3. Results

### 3.1. Species - and site-level characteristics did not outperform taxonomy in explaining variation in collision-mortality rates

The fitted data in the trait and taxonomic models included 2,212 records from 330 species, including 124 species known to occur in Europe. These records were compiled from 81 sources spanning 69 locations (Fig. S3). 63% of the fitted records were categorised as ‘high’ or ‘medium’ quality (meaning that group- or species-specific corrections for detectability were applied; see Thaxter et al. (2017)), and 37% of the records as ‘low’ quality (some corrections applied). The fitted data did not include any study whose quality was classified as ‘very low’ (i.e., no study where no form of detectability correction was applied).

We found that collision-mortality rate was positively associated with turbine capacity (trait model, Fig. S7: estimated slope: 1.1; 95% CI: 0.74-1.4; taxonomic model: 1.1 [0.80;1.4]). However, we did not find any association between species-level characteristics and collision-mortality rate (Fig. S7). Although low study quality had a negative effect on estimated rates, there was no significant difference between the effects of medium and high-quality studies. Further, model comparisons (Table S2) showed that the different fitted models did not differ from each other in terms of predictive performance, even when considering landscape predictors and despite negative associations of both terrain roughness (estimate: -0.53[-0.97;-0.10]) and elevation (-0.33[-0.58;-0.09]) with collision-mortality rates (note that roughness and elevation were positively correlated with Pearson’s r=0.55). In the trait model, the fixed effects explained a small proportion of the overall variation in collision mortality, while the random effects (including taxonomy) explained an important part of the variation (marginal pseudo-*R*^2^: 0.064; conditional pseudo-*R*^2^: 0.96). In the taxonomic model, the variation explained by the fixed effects was higher (marginal pseudo-*R*^2^: 0.53; conditional pseudo-*R*^2^: 0.96).

Given that species- and site-level characteristics did not explain an important part of the variation in collision-mortality rates and given that the trait models did not outperform the taxonomic model, we retained the taxonomic model for all further analyses, and we did not make inferences for additional European species based on trait values (Fig. S1); the taxonomic model was therefore employed to estimate the impact index at the species level (Fig. S8).

### 3.2. Risk estimation and patterns of risk

Among the 124 European species that figured in the fitted data, 108 had enough information to quantify vulnerability and therefore risk (Fig. S8). Across species, we found that the vulnerability and impact indices were negatively correlated (Pearson’s r: -0.36; p-value<0.001; Fig. 3), with average impacts decreasing significantly with increasing vulnerability. The species with the highest impact index (i.e. total estimated fatalities) was *Emberiza calandra* (corn bunting), while the species with the highest vulnerability index was *Uria aalge* (common murre). Median vulnerability and impact differed across taxonomic Orders (Fig. S9). The distribution of the impact index was right-skewed (Fig. S9), with many species exhibiting lower relative impacts and few species exhibiting higher relative impacts. Using 75% quantiles as thresholds (overlapping top 25% species in both dimensions), we found 3 species to be at high risk (Fig. 3), and 24 species to be at high latent risk. 81 species were classified as potential adapters or potential persisters. Using 50% quantiles meant an additional 9 species were considered at high risk, and an additional 18 species were considered at high latent risk (54 species in total at high (latent) risk when using 50% quantile thresholds).

**Fig. 3.**
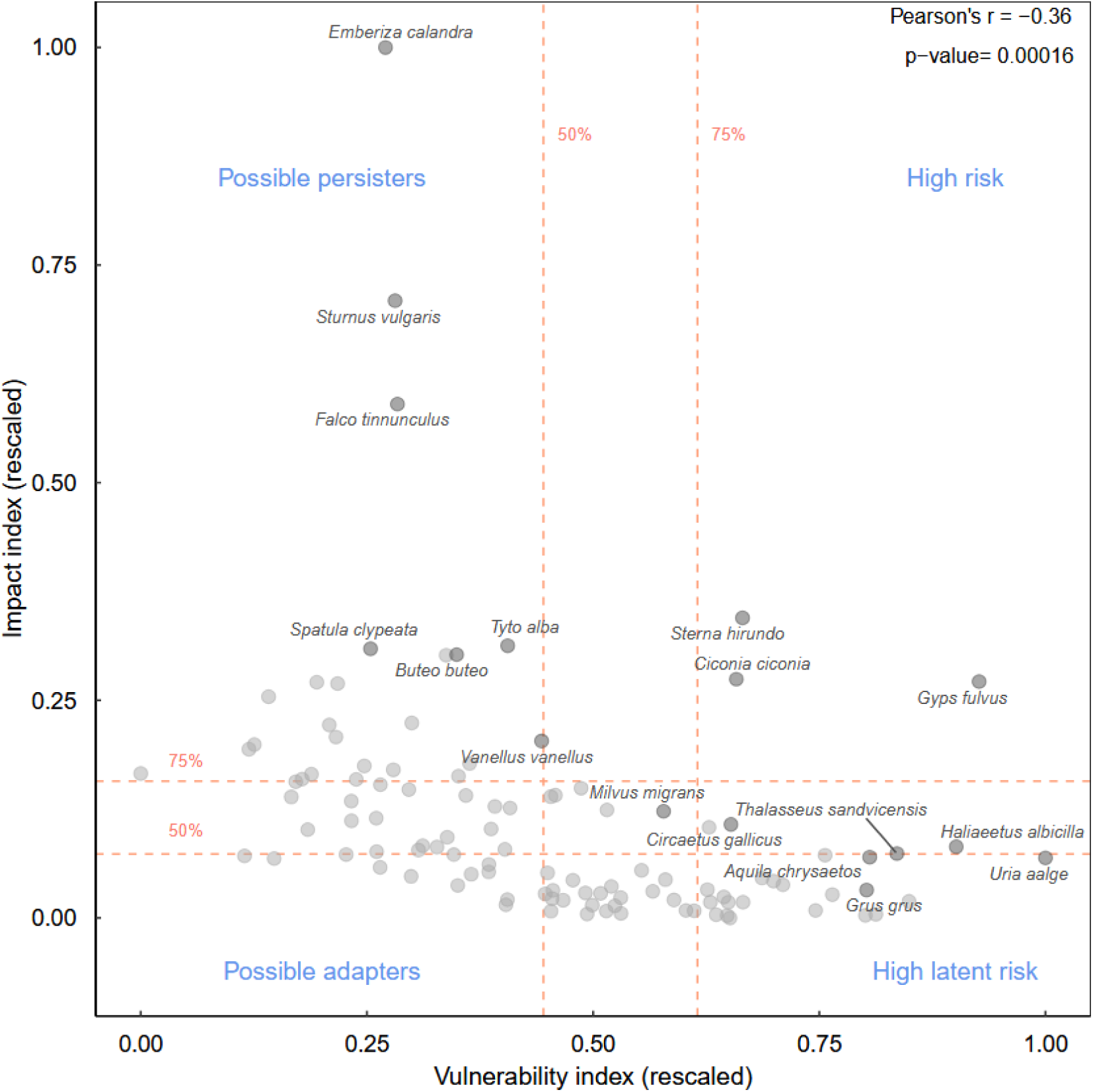
Estimated impact index against vulnerability index, and classification into risk categories, for 108 species that figured in the fatality-count data. Dashed lines represent 75% and 50% quantiles of the distributions for each axis of risk. We highlight some of the species in the data, but not all, for readability.

Plotting pseudo-species richness in the different risk categories highlighted hotspots of risk in Europe (Fig. S10). Parts of the Iberian Peninsula, of Northern Europe, and of the Balkans notably tended to concentrate more species at high (latent) risk. Using 50% quantiles for the classification of species into different risk categories did not affect the spatial patterns overall (Fig. S9; Fig. S10), although the number of species considered at high risk and high latent risk increased, therefore also expanding areas concentrating more species at higher risk into other regions. Combining information on wind turbine locations with the pseudo-species richness in the different risk categories, we obtained bivariate plots (Fig. 4; Fig. S12) showing regions which had both a higher density of turbines and a higher number of species at high risk or high latent risk, highlighting possible areas of conflict. Overall, the current turbine deployment had a higher degree of overlap with suitable habitats of species at lower relative vulnerability than with those of species at higher relative vulnerability (Fig. S13). For instance, turbine density was lower around central areas of the Iberian Peninsula, which also concentrated some of the species at higher risk (e.g. the Griffon vulture, *Gyps fulvus*). On the other hand, Germany tended to harbour a higher number of species at lower relative vulnerability and exhibited a higher density of turbines, but fewer species at higher relative vulnerability.

**Fig. 4.**
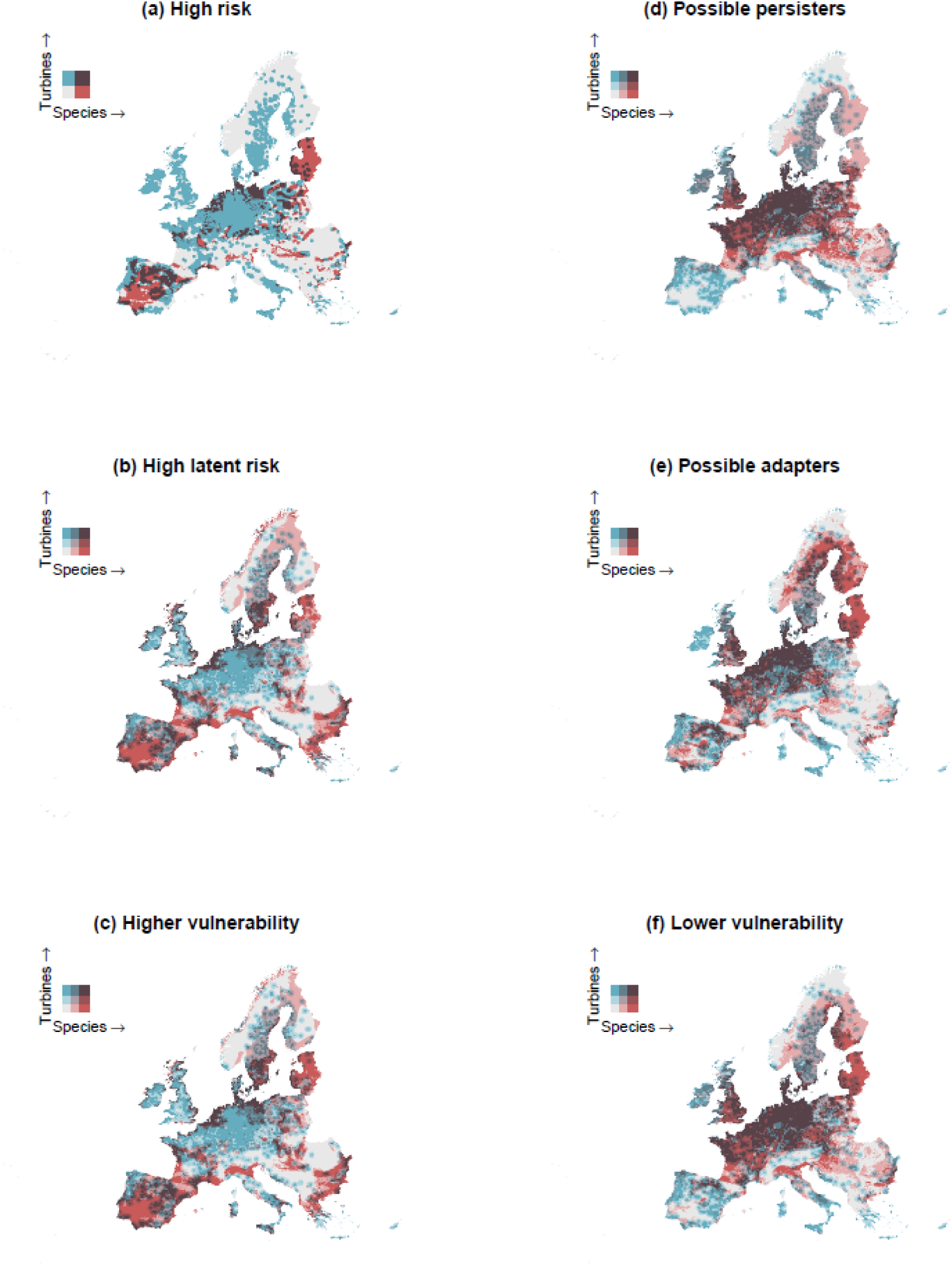
Spatial overlap between suitable habitats for species in the different risk categories or vulnerability groupings and current wind-turbine locations; (a) & (b) high risk and high latent risk species; (c) higher relative vulnerability (i.e., high risk and high latent risk species considered together); (d) & (e) possible persisters and possible adapters; (f) lower relative vulnerability (i.e., possible persisters and possible adapters considered together). Red areas harbour a larger number of species in a given risk category; blue areas harbour a larger number of turbines; purple areas indicate overlaps between areas of higher pseudo-species richness and higher turbine density. Here, risk categories were derived using the 75% quantiles of the distribution of the vulnerability and impact indices as thresholds. Note that we used only two breaks in (a) because no more than 3 high-risk species occurred simultaneously in any given grid cell, while most often only 2 species occurred.

## 4. Discussion

### 4.1. Key results and interpretations

Here, using a risk assessment approach built upon two dimensions (impact and vulnerability), we evaluated the current risk to 108 European birds from collisions with wind-energy facilities. Our estimation of the impact index relied on fatality counts which can be biased (e.g. with more detectable species more frequently reported; Nilsson et al. (2023)). Therefore, we emphasize that our results reflected the ‘reported’ risk; true risk might be underestimated for species for which fatalities remain underreported. However, although data biases may influence our results, all fatality records we considered were corrected for detectability in some way, with over 60% of the records including species- or group-specific corrections (Thaxter et al., 2017), and we found no evidence of further bias in the data (S10). This suggests that our results were robust to potential data biases. Further, the vulnerability index was estimated independently from the impact index, so the relative vulnerability groupings were unaffected by possible biases in the fatality-count data. Our study therefore provides a proof-of-concept for applying a risk framework to wind-turbine collisions at macroecological scales, which future studies can build upon, refine further and validate.

We found that reported risk was unevenly distributed in Europe, with risk hotspots in the Iberian Peninsula, in Northern Europe, in the Balkans, and in parts of Western and central Europe. The overlap between wind-turbine locations and species at higher relative vulnerability highlighted regions where conflicts between wind-energy facilities and bird conservation might currently arise. Our results however showed that the current deployment of wind turbines overlapped more with the suitable areas of species at lower relative risk than with that of species at higher relative risk, signalling that to now, turbine deployment may have avoided some areas harbouring relatively more vulnerable species.

Of the 108 considered species, we found 27 species to be at higher relative vulnerability, of which 6 belonged to the *Accipitriformes* Order (birds of prey), and 12 belonged to the *Charadriiformes* Order (shorebirds), when using a more conservative threshold value for the groupings (75% quantiles). Most species classified as possible persisters (i.e., higher relative impacts, but lower relative vulnerability) belonged to the *Passeriformes* Order (perching birds), suggesting that such species possibly sustain high impacts on their populations. Altogether, this highlights the effects of wind-energy developments on such bird Orders, in line with previous studies (Desholm, 2009; Erickson et al., 2014; Estellés-Domingo & López-López, 2024; Thaxter et al., 2017).

Unlike Thaxter et al. (2017), we found that the interspecific variation in collision-mortality rates could not be attributed to the ecological characteristics considered (but Thaxter et al. (2017) considered categorised dispersal distances, i.e, dispersal bands, whereas we used a continuous proxy of dispersal ability). Further, species- and site-level characteristics did not outperform taxonomy in estimating collision-mortality rates; and most of the variation explained in the different models was attributable to taxonomy. It could be that collision mortality is influenced by factors that we did not capture here, and possibly challenging to account for at large scales. For instance, weather, species behavioural plasticity, flight height, or fine-scale landscape variables (e.g., distance to breeding grounds) were difficult to capture in a quantitative synthesis such as ours but likely play an important role in collision mortality; it could also be that a high degree of context specificity hampers our ability to detect some effects.

Finally, species face multiple anthropogenic pressures. In a study of mortality causes for large birds over the Eurasian-African flyway, Serratosa et al. (2024) showed that mortality events related to energy infrastructure represented 49% of human-induced mortality events, but only a small proportion of these were attributed to collisions with wind-energy facilities; electrocutions and collisions with power lines represented most mortality events. In vultures, direct and indirect poisoning is a major source of human-induced mortality, and vulture protection may require curbing poisoning as well as collision mortality (Posillico et al., 2023; Sanz-Aguilar et al., 2015). Thus, as species face multiple pressures, ensuring that the risk posed by wind-energy facilities does not exacerbate the threats on populations is important to their long-term viability.

### 4.2. Possible applications

Our maps could be employed in decision-making tools, e.g. to constrain future scenarios of wind-power deployment at the European scale in energy-systems modelling, to improve biodiversity-inclusive spatial planning, to mitigate potential adverse impacts on birds, or to assess possible areas of conflicts and biodiversity impacts of different deployment scenarios. Our outputs could also be combined with other assessments of wind-power impacts (e.g. land-use change, noise, shadow flicker; McKenna et al., (2025)) for exploring spatial trade-offs and synergies of different impacts (social & ecological). However, we caution that our mapped outputs are not appropriate for fine-scale downscaling or local prioritisations, given that we could not capture some of the likely context specificity.

### 4.3. Research priorities

The amount of readily available collision-mortality data was an important limiting factor. We relied on data previously compiled by Thaxter et al. (2017). While there are recent initiatives aiming at centralising fatality records (e.g., see https://oscars-project.eu/projects/risky-wildlife-mortality-energy-and-transport-infrastructure; https://spbt.gr/en/database_victims_wind_turbines/), there is currently no protocol at the European scale for the collection and processing of such data. Unravelling the factors that influence collision risk at large scales may require developing frameworks for systematic processing of such data, which are currently scattered across studies and technical reports, implying important and possibly duplicated collection efforts for use in quantitative syntheses. Such systematically processed data, possibly enhanced with information on local conditions, would be valuable not only to understand what shapes collision mortality at large scales, but also to inform mitigation strategies and spatial planning. Standardised protocols for data collection would also enhance comparability across studies (Conkling et al., 2021) and help reduce potential sampling biases in quantitative synthesis such as ours. Systematically processed data could allow to further investigate seasonal, spatial or intraspecific variation in collision-mortality rates (which we did not consider here; our analyses considered species-level yearly averages, with data limitations also precluding the inclusion of interactions in the models). Thus, additional data could allow for further validation of our results and further characterisation of the inter- and intra-specific variation in collision mortality.

Finally, an important question is how collision mortality may impact species populations and demographic trends over large spatiotemporal scales (Duriez et al., 2023; May et al., 2019), and which collision-mortality rates we can consider as “acceptable” thresholds for the long-term viability of populations. Investigating such a question requires using mechanistic demographic models, which rely on demographic information that can be challenging to obtain across many species. By adapting a risk framework here which used readily available information across many species, we were able to assess which species were at higher relative risk; our approach, however, could not estimate an acceptable level of risk within a species.

### 4.4. Conclusion

Our work highlights the uneven spatial and taxonomic distribution of risk and can help identify areas of possible higher risk at the European scale. As more wind-energy facilities are deployed in the coming decades, mitigating the risk they pose to birds, particularly to species at higher risk, may be important for the long-term viability of the species.

## References

Amatulli, G., Domisch, S., Tuanmu, M.-N., Parmentier, B., Ranipeta, A., Malczyk, J., & Jetz, W. (2018). A suite of global, cross-scale topographic variables for environmental and biodiversity modeling. Scientific Data, 5(1), Article 1. 10.1038/sdata.2018.40

Bellard, C., Marino, C., Butt, N., Fernández-Palacios, J. M., Rigal, F., Robuchon, M., Lenoir, J., Irl, S., Benítez-López, A., Capdevila, P., Zhu, G., Caetano, G., Denelle, P., Philippe-Lesaffre, M., Schipper, A. M., Foden, W., Kissling, W. D., & Leclerc, C. (2024). A framework to quantify the vulnerability of insular biota to global change. https://hal.science/hal-04550966

Bellebaum, J., Korner-Nievergelt, F., Dürr, T., & Mammen, U. (2013). Wind turbine fatalities approach a level of concern in a raptor population. Journal for Nature Conservation, 21(6), 394–400. 10.1016/j.jnc.2013.06.001

Bernardino, J., Bispo, R., Costa, H., & Mascarenhas, M. (2013). Estimating bird and bat fatality at wind farms: A practical overview of estimators, their assumptions and limitations. New Zealand Journal of Zoology, 40(1), 63–74. 10.1080/03014223.2012.758155

Bird, J. P., Martin, R., Akçakaya, H. R., Gilroy, J., Burfield, I. J., Garnett, S. T., Symes, A., Taylor, J., Şekercioğlu, Ç. H., & Butchart, S. H. M. (2020). Generation lengths of the world’s birds and their implications for extinction risk. Conservation Biology, 34(5), 1252–1261. 10.1111/cobi.13486

Bürkner, P.-C. (2017). brms: An R Package for Bayesian Multilevel Models Using Stan. Journal of Statistical Software, 80, 1–28. 10.18637/jss.v080.i01

Bürkner, P.-C. (2018). Advanced Bayesian Multilevel Modeling with the R Package brms. The R Journal, 10(1), 395–411.

Bürkner, P.-C. (2021). Bayesian Item Response Modeling in R with brms and Stan. Journal of Statistical Software, 100, 1–54. 10.18637/jss.v100.i05

Cabrera-Cruz, S. A., & Villegas-Patraca, R. (2016). Response of migrating raptors to an increasing number of wind farms. Journal of Applied Ecology, 53(6), 1667–1675. 10.1111/1365-2664.12673

Carrete, M., Sánchez-Zapata, J. A., Benítez, J. R., Lobón, M., & Donázar, J. A. (2009). Large scale risk-assessment of wind-farms on population viability of a globally endangered long-lived raptor. Biological Conservation, 142(12), 2954–2961. 10.1016/j.biocon.2009.07.027

Carrete, M., Sánchez-Zapata, J. A., Benítez, J. R., Lobón, M., Montoya, F., & Donázar, J. A. (2012). Mortality at wind-farms is positively related to large-scale distribution and aggregation in griffon vultures. Biological Conservation, 145(1), 102–108. 10.1016/j.biocon.2011.10.017

Chapman, B. B., Brönmark, C., Nilsson, J.-Å., & Hansson, L.-A. (2011). The ecology and evolution of partial migration. Oikos, 120(12), 1764–1775. 10.1111/j.1600-0706.2011.20131.x

Chapman, M., Jung, M., Leclère, D., Boettiger, C., D.Augustynczik, A. L., Gusti, M., Ringwald, L., & Visconti, P. (2023). Meeting European conservation and restoration targets under future land-use demands. OSF. 10.31219/osf.io/ynqfx

Conkling, T. J., Loss, S. R., Diffendorfer, J. E., Duerr, A. E., & Katzner, T. E. (2021). Limitations, lack of standardization, and recommended best practices in studies of renewable energy effects on birds and bats. Conservation Biology, 35(1), 64–76. 10.1111/cobi.13457

Dahl, E. L., May, R., Hoel, P. L., Bevanger, K., Pedersen, H. C., Røskaft, E., & Stokke, B. G. (2013). White-tailed eagles (Haliaeetus albicilla) at the Smøla wind-power plant, Central Norway, lack behavioral flight responses to wind turbines. Wildlife Society Bulletin, 37(1), 66–74. 10.1002/wsb.258

Desholm, M. (2009). Avian sensitivity to mortality: Prioritising migratory bird species for assessment at proposed wind farms. Journal of Environmental Management, 90(8), 2672–2679. 10.1016/j.jenvman.2009.02.005

Directorate-General for Environment (European Commission), EuroCARE GmbH Bonn, IIASA, & UNEP-WCMC. (2024). *BIOCLIMA: Assessing land use, climate and biodiversity impacts of land based climate mitigation and biodiversity policies in the EU : technical report*. Publications Office of the European Union. https://data.europa.eu/doi/10.2779/37660

Domínguez del Valle, J., Cervantes Peralta, F., & Jaquero Arjona, M. I. (2020). Factors affecting carcass detection at wind farms using dogs and human searchers. Journal of Applied Ecology, 57(10), 1926–1935. 10.1111/1365-2664.13714

Drewitt, A. L., & Langston, R. H. W. (2008). Collision Effects of Wind-power Generators and Other Obstacles on Birds. Annals of the New York Academy of Sciences, 1134(1), 233–266. 10.1196/annals.1439.015

Duriez, O., Pilard, P., Saulnier, N., Boudarel, P., & Besnard, A. (2023). Windfarm collisions in medium-sized raptors: Even increasing populations can suffer strong demographic impacts. Animal Conservation, 26(2), 264–275. 10.1111/acv.12818

Erickson, W. P., Wolfe, M. M., Bay, K. J., Johnson, D. H., & Gehring, J. L. (2014). A Comprehensive Analysis of Small-Passerine Fatalities from Collision with Turbines at Wind Energy Facilities. PLOS ONE, 9(9), e107491. 10.1371/journal.pone.0107491

Estellés-Domingo, I., & López-López, P. (2024). Effects of wind farms on raptors: A systematic review of the current knowledge and the potential solutions to mitigate negative impacts. Animal Conservation. 10.1111/acv.12988

Farfán, M. Á., Díaz-Ruiz, F., Duarte, J., Martín-Taboada, A., & Muñoz, A.-R. (2023). Wind farms and Griffon Vultures: Evidence that under certain conditions history is not-always turbulent. Global Ecology and Conservation, 48, e02728. 10.1016/j.gecco.2023.e02728

Fernández-Bellon, D., Wilson, M. W., Irwin, S., & O’Halloran, J. (2019). Effects of development of wind energy and associated changes in land use on bird densities in upland areas. Conservation Biology, 33(2), 413–422. 10.1111/cobi.13239

Fischereit, J., Olsen, B.T., Imberger, M., Vedel, H., Hahmann, A.N., Larsén, X. G.: (n.d.). Modelling wind farm effects in HARMONIE-AROME – part 2: Wind turbine database and application to Europe, Geoscientific Model Development. In preparation.

Foden, W. B., Butchart, S. H. M., Stuart, S. N., Vié, J.-C., Akçakaya, H. R., Angulo, A., DeVantier, L. M., Gutsche, A., Turak, E., Cao, L., Donner, S. D., Katariya, V., Bernard, R., Holland, R. A., Hughes, A. F., O’Hanlon, S. E., Garnett, S. T., Şekercioğlu, Ç. H., & Mace, G. M. (2013). Identifying the World’s Most Climate Change Vulnerable Species: A Systematic Trait-Based Assessment of all Birds, Amphibians and Corals. PLOS ONE, 8(6), e65427. 10.1371/journal.pone.0065427

Foden, W. B., Young, B. E., Akçakaya, H. R., Garcia, R. A., Hoffmann, A. A., Stein, B. A., Thomas, C. D., Wheatley, C. J., Bickford, D., Carr, J. A., Hole, D. G., Martin, T. G., Pacifici, M., Pearce-Higgins, J. W., Platts, P. J., Visconti, P., Watson, J. E. M., & Huntley, B. (2019). Climate change vulnerability assessment of species. WIREs Climate Change, 10(1), e551. 10.1002/wcc.551

Gómez-Catasús, J., Garza, V., & Traba, J. (2018). Wind farms affect the occurrence, abundance and population trends of small passerine birds: The case of the Dupont’s lark. Journal of Applied Ecology, 55(4), 2033–2042. 10.1111/1365-2664.13107

Harrison, X. A. (2014). Using observation-level random effects to model overdispersion in count data in ecology and evolution. PeerJ, 2, e616. 10.7717/peerj.616

Huso, M. M. P., & Dalthorp, D. (2014). Accounting for unsearched areas in estimating wind turbine-caused fatality. The Journal of Wildlife Management, 78(2), 347–358. 10.1002/jwmg.663

Hyman, A. A., Crone, E. R., Benson, A., Dunham, J., Lynch, A. J., Thompson, L., & Mims, M. C. (2025). Exposure, sensitivity, or adaptive capacity? Reviewing assessments that use only two of three elements of climate change vulnerability. Conservation Science and Practice, 7(1), e13293. 10.1111/csp2.13293

Intergovernmental Panel on Climate Change. (2007). *Climate change 2007: Impacts, adaptation and vulnerability. Contribution of working group II to the fourth assessment report of the Intergovernmental Panel on Climate Change*, 2007. Cambridge, England: Cambridge University Press.

Intergovernmental Panel on Climate Change. (2014). *Climate change 2014: Impacts, adaptation, and vulnerability. Contributions of working group II to the fifth assessment report*. Cambridge, England/New York, NY: Cambridge University Press.

Jung, M. (2023). An integrated species distribution modelling framework for heterogeneous biodiversity data. Ecological Informatics, 76, 102127. 10.1016/j.ecoinf.2023.102127

Kuvlesky Jr., W. P., Brennan, L. A., Morrison, M. L., Boydston, K. K., Ballard, B. M., & Bryant, F. C. (2010). Wind Energy Development and Wildlife Conservation: Challenges and Opportunities. The Journal of Wildlife Management, 71(8), 2487–2498. 10.2193/2007-248

Loss, S. R., Will, T., & Marra, P. P. (2013). Estimates of bird collision mortality at wind facilities in the contiguous United States. Biological Conservation, 168, 201–209. 10.1016/j.biocon.2013.10.007

Marques, A. T., Batalha, H., Rodrigues, S., Costa, H., Pereira, M. J. R., Fonseca, C., Mascarenhas, M., & Bernardino, J. (2014). Understanding bird collisions at wind farms: An updated review on the causes and possible mitigation strategies. Biological Conservation, 179, 40–52. 10.1016/j.biocon.2014.08.017

Marques, A. T., Santos, C. D., Hanssen, F., Muñoz, A.-R., Onrubia, A., Wikelski, M., Moreira, F., Palmeirim, J. M., & Silva, J. P. (2020). Wind turbines cause functional habitat loss for migratory soaring birds. Journal of Animal Ecology, 89(1), 93–103. 10.1111/1365-2656.12961

May, R., Masden, E. A., Bennet, F., & Perron, M. (2019). Considerations for upscaling individual effects of wind energy development towards population-level impacts on wildlife. Journal of Environmental Management, 230, 84–93. 10.1016/j.jenvman.2018.09.062

May, R., Reitan, O., Bevanger, K., Lorentsen, S.-H., & Nygård, T. (2015). Mitigating wind-turbine induced avian mortality: Sensory, aerodynamic and cognitive constraints and options. Renewable and Sustainable Energy Reviews, 42, 170–181. 10.1016/j.rser.2014.10.002

McKenna, R., Lilliestam, J., Heinrichs, H. U., Weinand, J., Schmidt, J., Staffell, I., Hahmann, A. N., Burgherr, P., Burdack, A., Bucha, M., Chen, R., Klingler, M., Lehmann, P., Lowitzsch, J., Novo, R., Price, J., Sacchi, R., Scherhaufer, P., Schöll, E. M., … Camargo, L. R. (2025). System impacts of wind energy developments: Key research challenges and opportunities. Joule, 9(1). 10.1016/j.joule.2024.11.016

Myhrvold, N. P., Baldridge, E., Chan, B., Sivam, D., Freeman, D. L., & Ernest, S. K. M. (2015). An amniote life-history database to perform comparative analyses with birds, mammals, and reptiles. Ecology, 96(11), 3109–3109. 10.1890/15-0846R.1

Nilsson, A. L. K., Molværsmyr, S., Breistøl, A., & Systad, G. H. R. (2023). Estimating mortality of small passerine birds colliding with wind turbines. Scientific Reports, 13(1), Article 1. 10.1038/s41598-023-46909-z

Pacifici, M., Santini, L., Marco, M. D., Baisero, D., Francucci, L., Marasini, G. G., Visconti, P., & Rondinini, C. (2013). Generation length for mammals. Nature Conservation, 5, 89–94. 10.3897/natureconservation.5.5734

Posillico, M., Costanzo, A., Bottoni, S., Altea, T., Opramolla, G., Pascazi, A., Panella, M., & Ambrosini, R. (2023). Reported mortality of Griffon Vulture Gyps fulvus in central Italy and indications for conservation and management. Bird Conservation International, 33, e68. 10.1017/S0959270923000199

Prener, C., Grossenbacher, T., & Zehr, A. (2022). biscale: Tools and Palettes for Bivariate Thematic Mapping (p. 1.0.0) [Dataset]. 10.32614/CRAN.package.biscale

R Core Team. (n.d.). R: A Language and Environment for Statistical Computing, R Foundation for Statistical Computing, Vienna, Austria. https://www.R-project.org/

Santangeli, A., Butchart, S. H. M., Pogson, M., Hastings, A., Smith, P., Girardello, M., & Moilanen, A. (2018). Mapping the global potential exposure of soaring birds to terrestrial wind energy expansion. Ornis Fennica, 95(1), Article 1. 10.51812/of.133925

Santos, C. D., Ramesh, H., Ferraz, R., Franco, A. M. A., & Wikelski, M. (2022a). Factors influencing wind turbine avoidance behaviour of a migrating soaring bird. Scientific Reports, 12(1), Article 1. 10.1038/s41598-022-10295-9

Santos, C. D., Ramesh, H., Ferraz, R., Franco, A. M. A., & Wikelski, M. (2022b). Factors influencing wind turbine avoidance behaviour of a migrating soaring bird. Scientific Reports, 12, 6441. 10.1038/s41598-022-10295-9

Sanz-Aguilar, A., Sánchez-Zapata, J. A., Carrete, M., Benítez, J. R., Ávila, E., Arenas, R., & Donázar, J. A. (2015). Action on multiple fronts, illegal poisoning and wind farm planning, is required to reverse the decline of the Egyptian vulture in southern Spain. Biological Conservation, 187, 10–18. 10.1016/j.biocon.2015.03.029

Schaub, M. (2012). Spatial distribution of wind turbines is crucial for the survival of red kite populations. Biological Conservation, 155, 111–118. 10.1016/j.biocon.2012.06.021

Schöll, E. M., & Nopp-Mayr, U. (2021). Impact of wind power plants on mammalian and avian wildlife species in shrub- and woodlands. Biological Conservation, 256, 109037. 10.1016/j.biocon.2021.109037

Serratosa, J., Oppel, S., Rotics, S., Santangeli, A., Butchart, S. H. M., Cano-Alonso, L. S., Tellería, J. L., Kemp, R., Nicholas, A., Kalvāns, A., Galarza, A., Franco, A. M. A., Andreotti, A., Kirschel, A. N. G., Ngari, A., Soutullo, A., Bermejo-Bermejo, A., Botha, A. J., Ferri, A., … Jones, V. R. (2024). Tracking data highlight the importance of human-induced mortality for large migratory birds at a flyway scale. Biological Conservation, 293, 110525. 10.1016/j.biocon.2024.110525

Sheard, C., Neate-Clegg, M. H. C., Alioravainen, N., Jones, S. E. I., Vincent, C., MacGregor, H. E. A., Bregman, T. P., Claramunt, S., & Tobias, J. A. (2020). Ecological drivers of global gradients in avian dispersal inferred from wing morphology. Nature Communications, 11(1), Article 1. 10.1038/s41467-020-16313-6

Shiomi, K. (2022). Possible link between brain size and flight mode in birds: Does soaring ease the energetic limitation of the brain? Evolution, 76(3), 649–657. 10.1111/evo.14425

Smallwood, K. S. (2013). Comparing bird and bat fatality-rate estimates among North American wind-energy projects. Wildlife Society Bulletin, 37(1), 19–33. 10.1002/wsb.260

Smeraldo, S., Bosso, L., Fraissinet, M., Bordignon, L., Brunelli, M., Ancillotto, L., & Russo, D. (2020). Modelling risks posed by wind turbines and power lines to soaring birds: The black stork (Ciconia nigra) in Italy as a case study. Biodiversity and Conservation, 29(6), 1959–1976. 10.1007/s10531-020-01961-3

Thaxter, C. B., Buchanan, G. M., Carr, J., Butchart, S. H. M., Newbold, T., Green, R. E., Tobias, J. A., Foden, W. B., O’Brien, S., & Pearce-Higgins, J. W. (2017). Bird and bat species’ global vulnerability to collision mortality at wind farms revealed through a trait-based assessment. Proceedings of the Royal Society B: Biological Sciences, 284(1862), 20170829. 10.1098/rspb.2017.0829

Tobias, J. A., Sheard, C., Pigot, A. L., Devenish, A. J. M., Yang, J., Sayol, F., Neate-Clegg, M. H. C., Alioravainen, N., Weeks, T. L., Barber, R. A., Walkden, P. A., MacGregor, H. E. A., Jones, S. E. I., Vincent, C., Phillips, A. G., Marples, N. M., Montaño-Centellas, F. A., Leandro-Silva, V., Claramunt, S., … Schleuning, M. (2022). AVONET: Morphological, ecological and geographical data for all birds. Ecology Letters, 25(3), 581–597. 10.1111/ele.13898

Vehtari, A., Gelman, A., & Gabry, J. (2017). Practical Bayesian model evaluation using leave-one-out cross-validation and WAIC. Statistics and Computing, 27(5), 1413–1432. 10.1007/s11222-016-9696-4

Voigt, C. C. (2021). Insect fatalities at wind turbines as biodiversity sinks. Conservation Science and Practice, 3(5), e366. 10.1111/csp2.366

Voigt, C. C., Straka, T. M., & Fritze, M. (2019). Producing wind energy at the cost of biodiversity: A stakeholder view on a green-green dilemma. Journal of Renewable and Sustainable Energy, 11(6), 063303. 10.1063/1.5118784

Vuichard, P., Broughel, A., Wüstenhagen, R., Tabi, A., & Knauf, J. (2022). Keep it local and bird-friendly: Exploring the social acceptance of wind energy in Switzerland, Estonia, and Ukraine. Energy Research & Social Science, 88, 102508. 10.1016/j.erss.2022.102508

Watanabe, Y. Y. (2016). Flight mode affects allometry of migration range in birds. Ecology Letters, 19(8), 907–914. 10.1111/ele.12627

Weeks, B. C., O’Brien, B. K., Chu, J. J., Claramunt, S., Sheard, C., & Tobias, J. A. (2022). Morphological adaptations linked to flight efficiency and aerial lifestyle determine natal dispersal distance in birds. Functional Ecology, 36(7), 1681–1689. 10.1111/1365-2435.14056

Wind Europe. (2022). Wind energy in Europe. 2022 Statistics and the outlook for 2023-2027. https://windeurope.org/intelligence-platform/product/wind-energy-in-europe-2022-statistics-and-the-outlook-for-2023-2027/

Zurell, D., Zimmermann, N. E., Gross, H., Baltensweiler, A., Sattler, T., & Wüest, R. O. (2020). Testing species assemblage predictions from stacked and joint species distribution models. Journal of Biogeography, 47(1), 101–113. 10.1111/jbi.13608

