## Supporting_Information for "Risk to European birds from collisions with wind-energy facilities"

#### S1. Conceptual framework and objectives

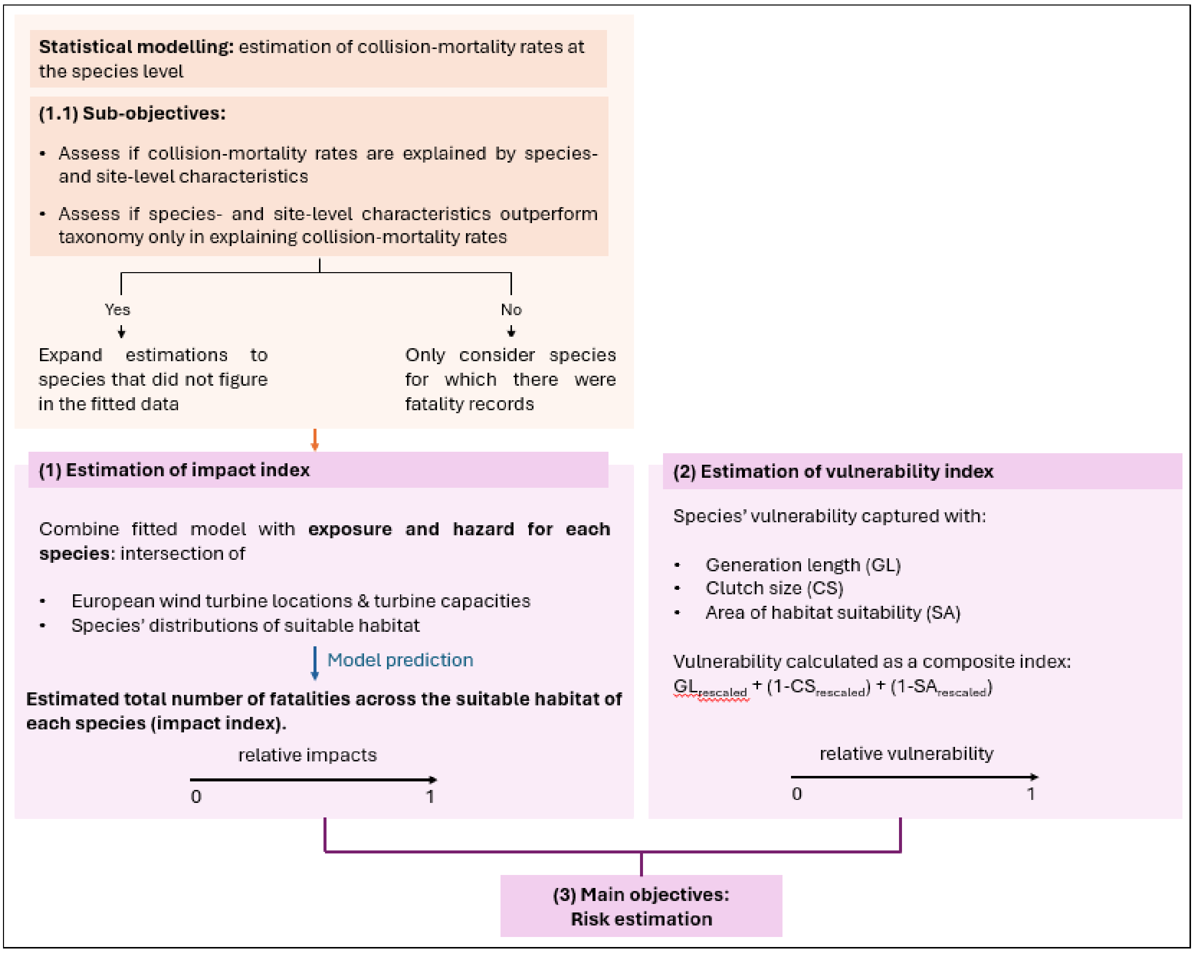


**Fig. S1. Workflow and aims.** We adapted a risk framework (Foden et al., 2019) to assess the relative, comparative risk to European bird species posed by collisions with wind-energy facilities. To determine the taxonomic scope of our study, we also assessed whether collision-mortality rates were associated with species- and site-level characteristics (Sub-objectives 1.1), therefore investigating if collision-mortality rates could be estimated for all European species based on such possible associations, and if we could thus extend the risk estimation to species for which there were no fatality records.

#### S2. From collision-mortality rates to fatality counts

To convert the standardised collision-mortality rates (number of fatalities per year-turbine) to fatality counts, we multiplied the standardised rates in the original data with study duration and number of turbines within each wind-energy facility. Since doing so did not necessarily generate integers – required to fit the models with a Poisson error distribution–, we rounded the estimates to the nearest integers. To get more precise estimates of the number of fatalities with a more accurate rounding, we increased the study duration by 10 years, multiplying both estimated rates and study duration by 10 before rounding. This allowed for a better estimation of rounded counts since initial counts of e.g., 0.2 over 1 year became 2 over 10 years, without affecting the rate itself. We further verified that the procedure did not affect the estimated rates, by comparing the initial standardised collision-mortality rates with collision-mortality rates derived from the rounded counts (Fig. S2).

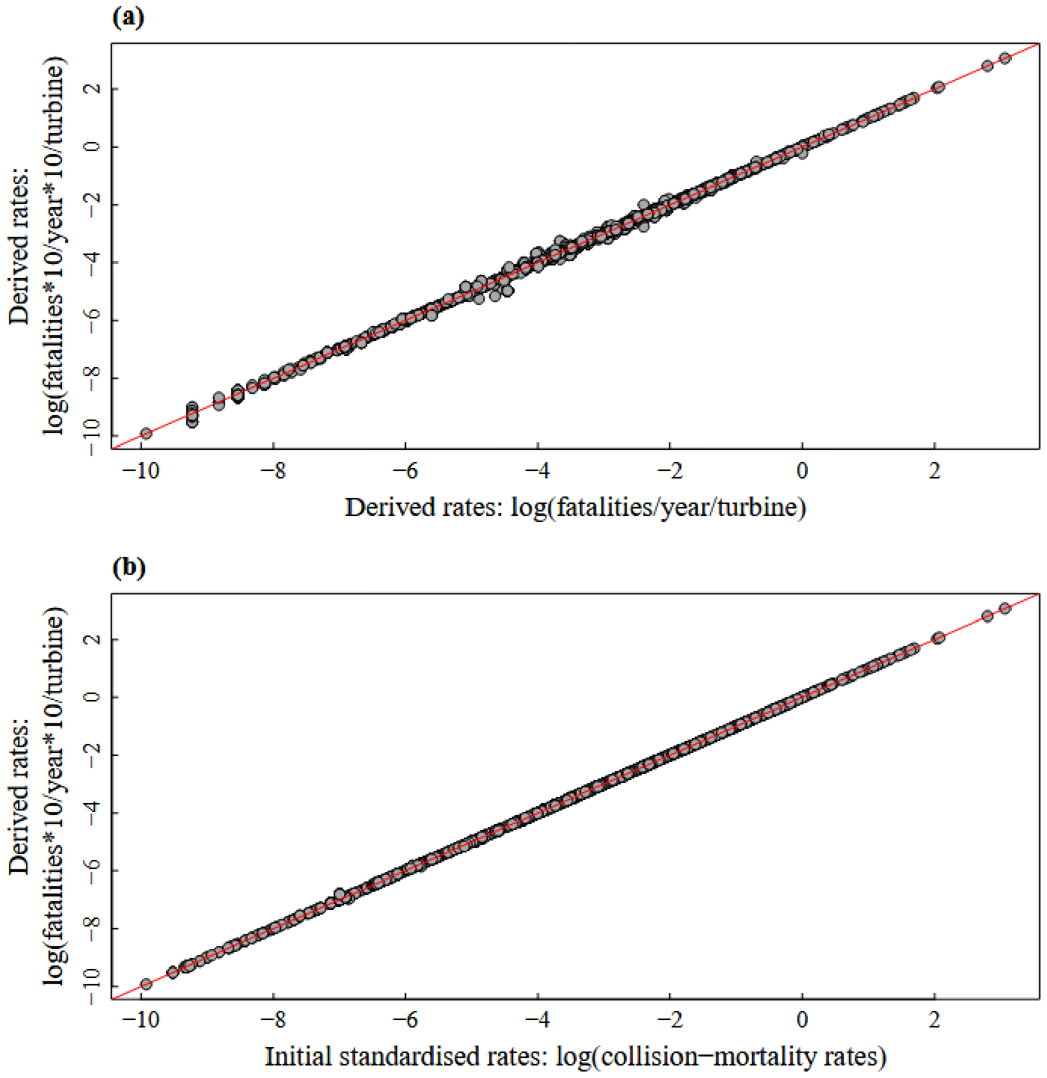


**Fig. S2.** Collision-mortality rates **(a)** derived from the original compiled data, calculated after converting the original standardised collision-mortality rates to fatality counts, either after multiplying original rates and study duration by 10 before rounding (y-axis), or rounding values directly without multiplying by 10 (x-axis). **(b)**: Derived collision-mortality rates we used (from counts and study duration multiplied by 10; y-axis) against original standardised collision-mortality rates from the compiled data.

#### S3. Fatality records

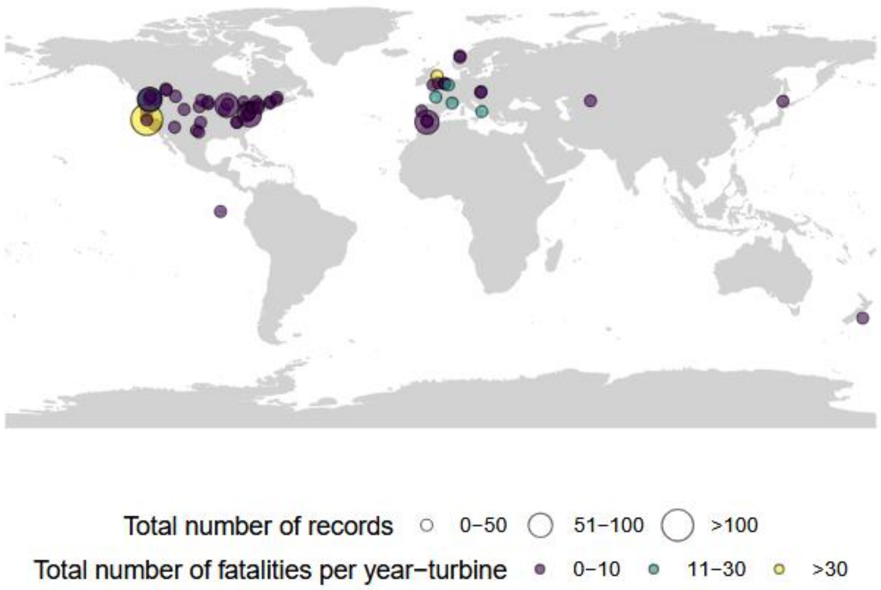


**Fig. S3.** Geographical location of the wind-energy facilities included in the fitted data (central longitude and central latitude of wind turbines within the facilities), number of records at each site (point size; in the data, each record provided fatality numbers for a given species), and total number of fatalities per year-turbine at each site across records (point colour). Land masses were obtained from the 'rnaturalearth' R package (Massicotte et al., 2023).

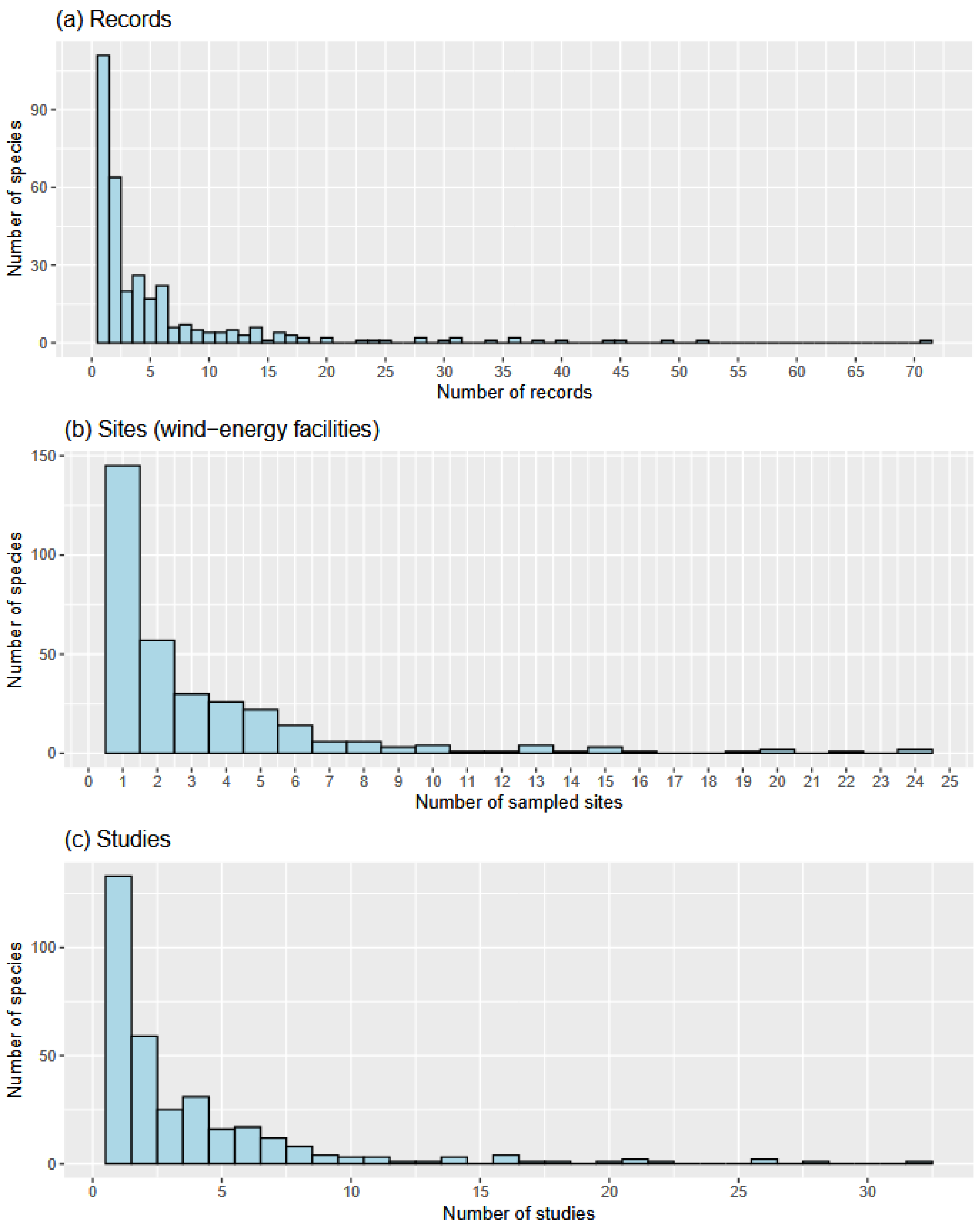


**Fig. S4.** Sampling of species: distribution of **(a)** number of records (in the fatality-count data, each record provided fatality numbers for a given species at a given site), **(b)** number of sites (wind-energy facilities) and **(c)** number of studies across sampled species.

#### S4. Buffer sizes

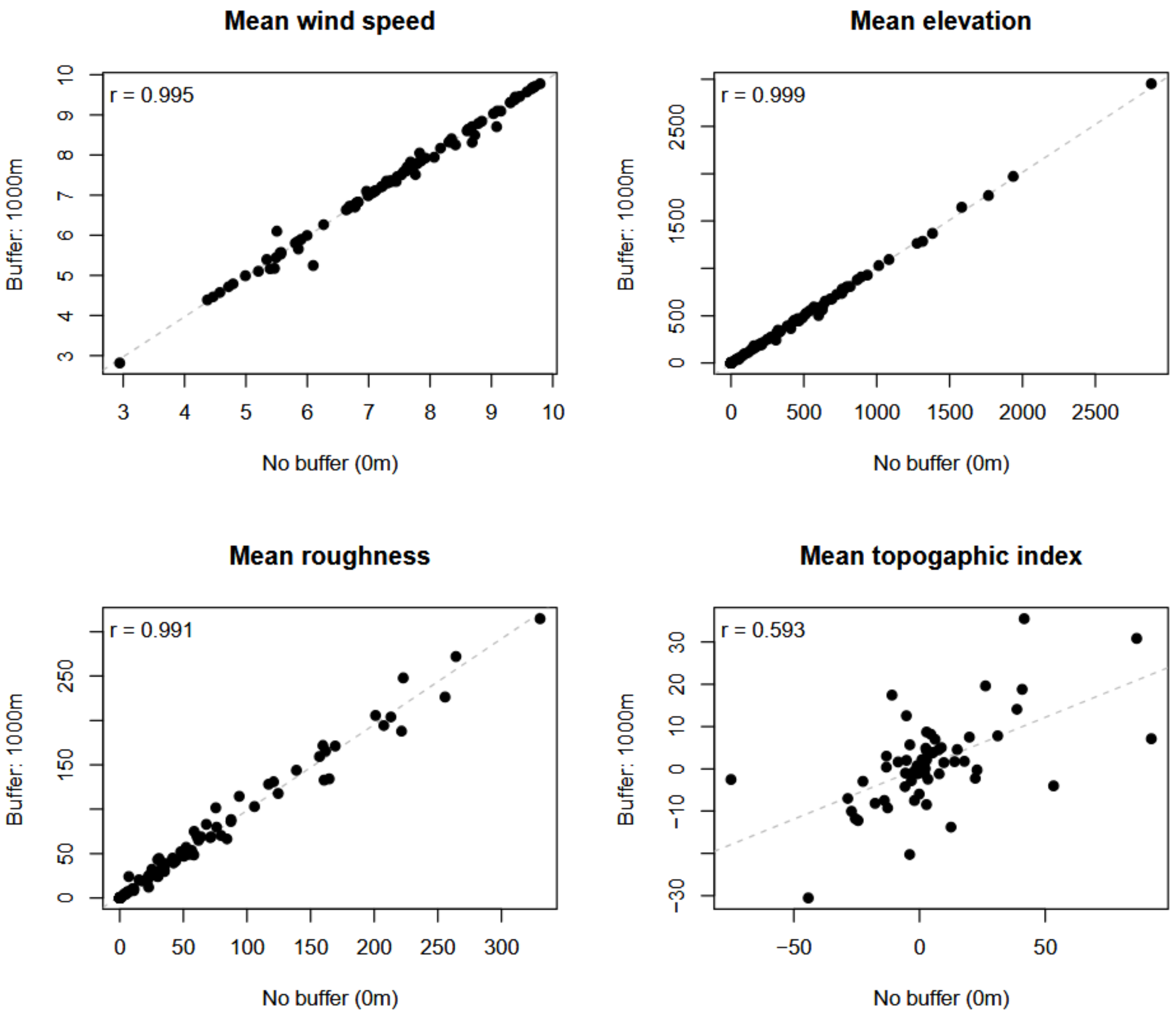


**Fig. S5.** Values of mean wind speed at a height of 100 metres (m/s), mean elevation (m), mean roughness (m), and mean topographic index at the central locations of the wind-energy facilities, extracted using two buffer sizes around the locations: 0 m (no buffer) and 1000 m. Each plot shows the values extracted with no buffer against the values extracted with a buffer of 1000 m. Pearson's correlation coefficients are reported in the top-left corners of the plots. The dotted grey lines are the lines of best fit given a linear model.

#### S5. Flight mode classification

We identified species that could be considered as soaring in the fatality-count data. The data spanned 330 species occurring in Europe and North America and encompassed 65 Families and 20 Orders. Following Santangeli et al. (2018), Shiomi (2022) and Watanabe (2016), we considered species in the Accipitriformes Order (22 species in the dataset), in the Falconiformes Order (7 species), in the Cathartiformes Order (1 species), and in the Pelecaniformes Order (10 species) as soaring. We additionally considered species in the Gruidae Family (2 species) and species in the Procellariidae Family (2 species). Out of 4 species occurring in the data, we included one species in the *Corvus* genus: *Corvus corax*. Finally, we considered the 18 species in the Laridae Family as soarers. We thus identified

65 species as soarers, out of 330 species. As emphasized by Shiomi (2022), birds use a combination of different flight modes. However, to assess whether mostly soaring species in particular had higher collision-mortality rates on average (Santangeli et al., 2018), we grouped species into two broad categories (soarers versus others), with soarers assumed to use soaring for a significantly larger proportion of their flight time than other species.

#### S6. Linearisation of the relationship between turbine capacity and fatality counts

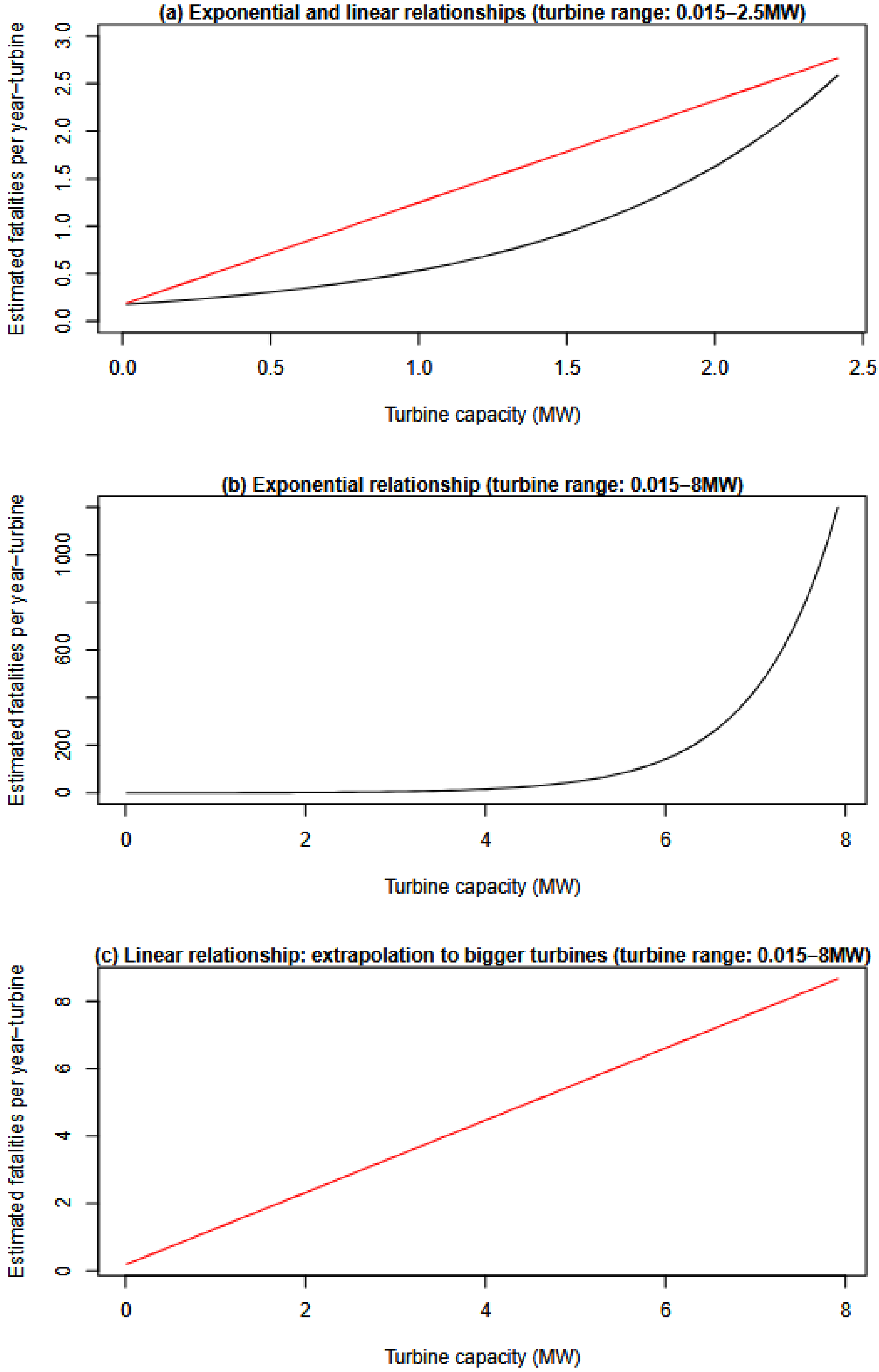


**Fig. S6.** Linearisation of the relationship between turbine capacity and fatality counts per year-turbine (e.g. collision-mortality rates). **(a)** Turbine capacity in the fitted data ranged between 0.015MW and

2.5MW, while turbines with a capacity of up to 8MW occurred in Europe. **(b)** Given that the models were not fitted on data encompassing turbines with a capacity larger than 2.5MW, assuming an exponential relationship would lead to unrealistic estimations of fatality numbers for large capacities. For each species, we used the model to predict values for capacities of 0.015MW and 2.5MW and estimated the slope assuming a linear relationship. **(c)** We then used the estimated slopes to predict fatality counts given any turbine capacity. The example shown here is based on estimates for *Gyps fulvus*.

#### S7. Vulnerability components

**Table S1.** Pearson's correlation coefficients among the different dimensions of the vulnerability index (CS: clutch size; GL: generation length; SA: area of habitat suitability). Note that SA was obtained by weighting the surface area of each grid cell by its suitability value and then summing weighted surface areas across all cells.

| | **$1\text{-}CS_{rescaled}$** | **$GL_{rescaled}$** | **$1\text{-}SA_{rescaled}$** |
|---|---|---|---|
| **$1\text{-}CS_{rescaled}$** | 1 | 0.49 | 0.17 |
| **$GL_{rescaled}$** | 0.49 | 1 | 0.29 |
| **$1\text{-}SA_{rescaled}$** | 0.17 | 0.29 | 1 |

#### S8. Fitted data and model outputs

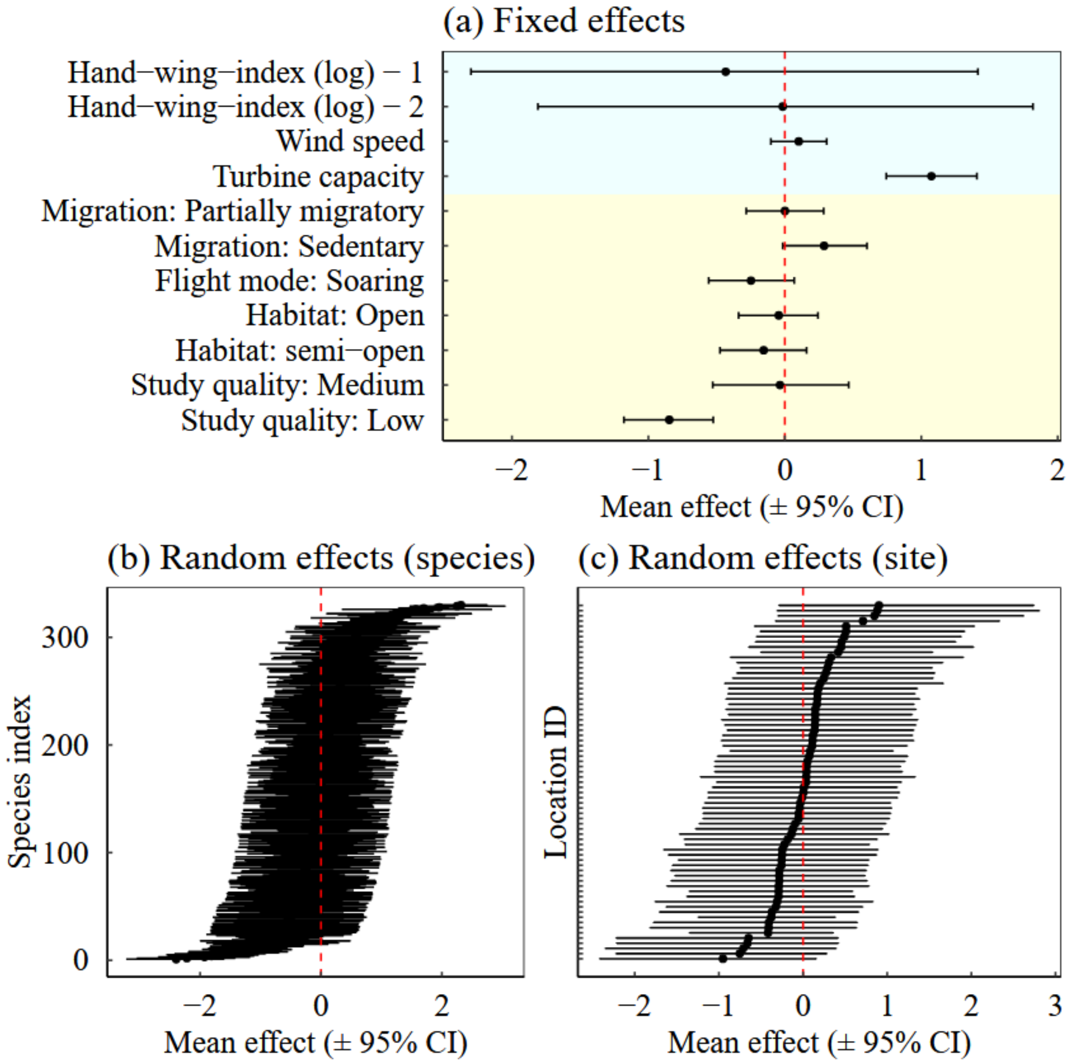


**Fig. S7.** (a) Estimated effects of the fixed predictors for the model fitted to explain collision-mortality rate by species- and site-level characteristics (‘trait’ model), mean ±95% credible interval, plotted on the log-scale. Light blue backgrounds highlight continuous predictors, while light yellow backgrounds highlight categorical predictors. Note that model estimates are presented on the log-scale. (b) Random-effects estimates of species identity. (c) Random-effects estimates of site (location) identity.

**Table S2.** Predictive accuracy differences (measured with expected log pointwise predictive density, termed "elpd" differences, among the alternative models tested). Larger elpd values tend to indicate better predictive performance; a model can be considered to have a better predictive performance than another when the elpd difference between the models exceeds 4, and when the standard error of the elpd estimate is small (Vehtari et al., 2017). Here, the model with the highest estimated elpd figures in the first row, and further models are ordered according to decreasing elpd differences. Given the relatively large values of the standard errors compared with the value of the elpd estimates, we concluded that the different models did not differ significantly from each other in terms of predictive performance. Thus, the trait model with highest elpd estimate did not significantly outperform the taxonomic model in terms of predictive performance.

| **Model** | **elpd difference** | **Standard error of elpd difference** |
|---|---|---|
| "Base" trait model with elevation added | 0 | 0 |
| **Taxonomic model** | **-0.8** | **6.1** |
| "Base" trait model with habitat openness swapped for habitat categories | -2.2 | 6.1 |
| "Base" trait model with terrain roughness added | -8.9 | 6.0 |
| "Base" trait model with topographic index added | -9.2 | 5.9 |
| "Base" trait model (considering habitat openness and mean wind speed) | -13.2 | 6.3 |

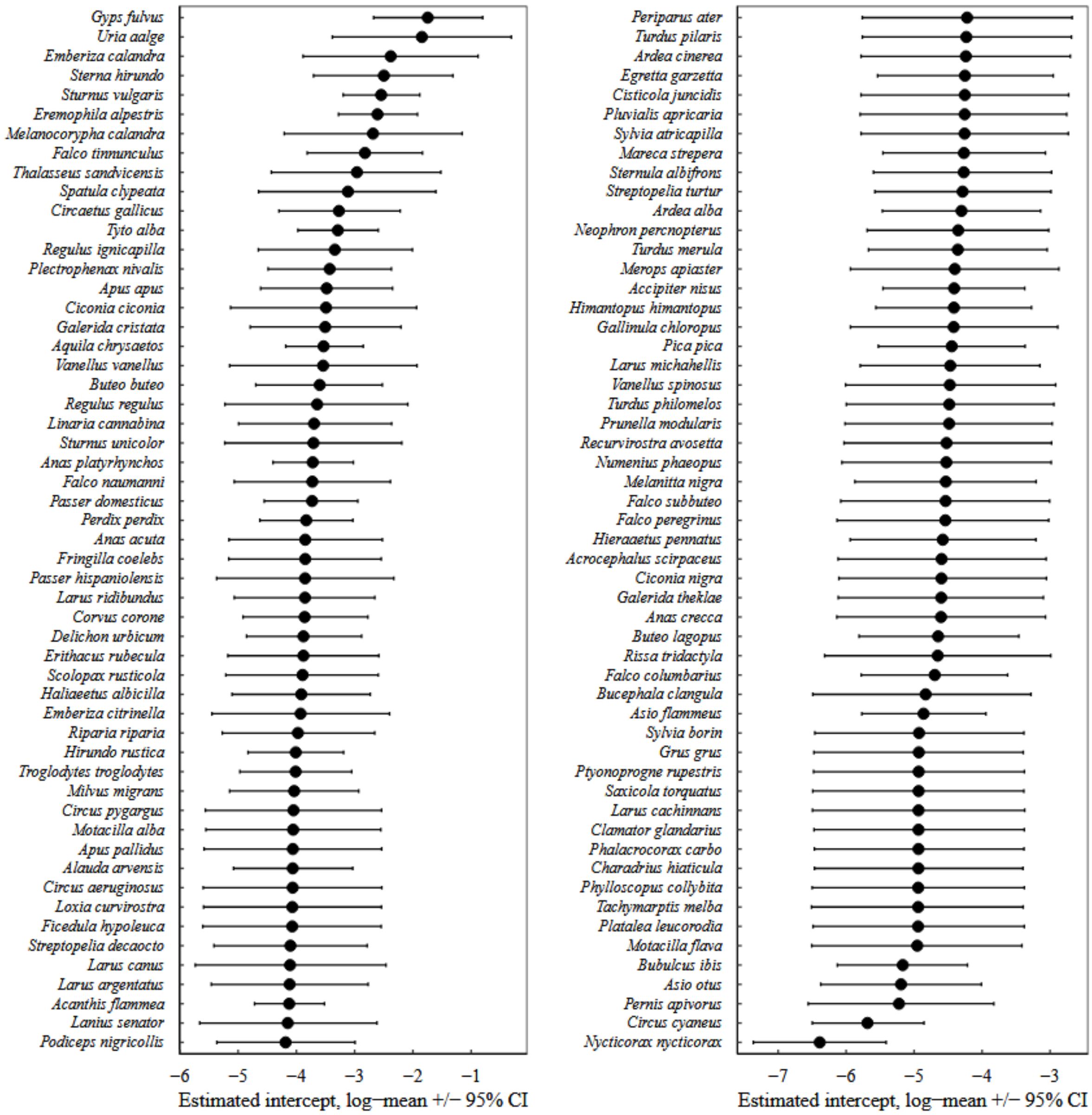


**Fig. S8.** Estimated intercepts from the taxonomic model (±95% credible interval) for the 108 species considered in the risk estimation.

### S9. Risk estimation & mapping

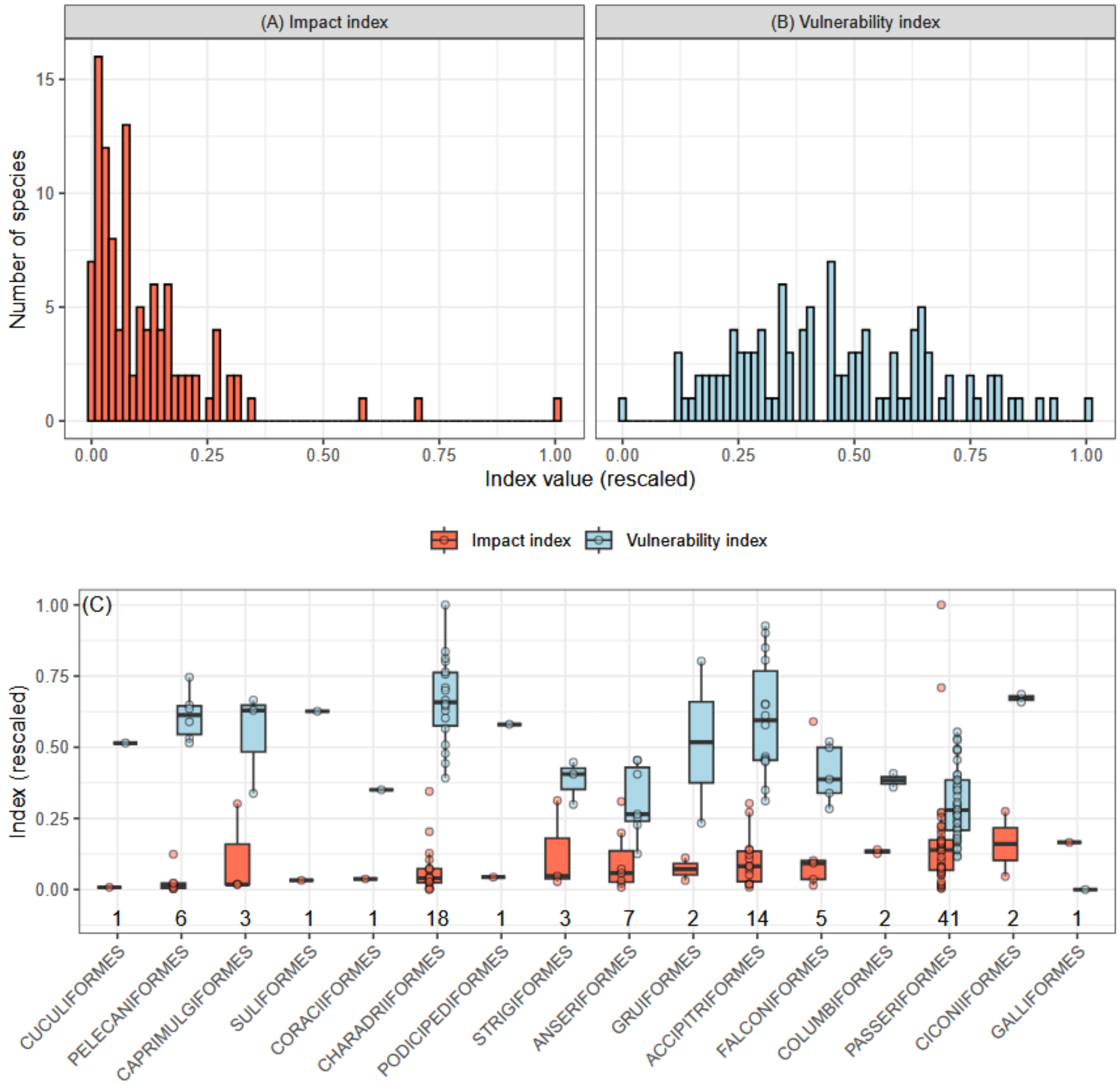


**Fig. S9.** Distribution of the impact index **(A)** and the vulnerability index **(B)** across the 108 species considered in the risk estimation. **(C)** Distribution of the indices by bird taxonomic Orders. Orders are sorted from left to right according to increasing value of median impact index. The numbers above the x-axis indicate the sample size (number of species) in each Order.

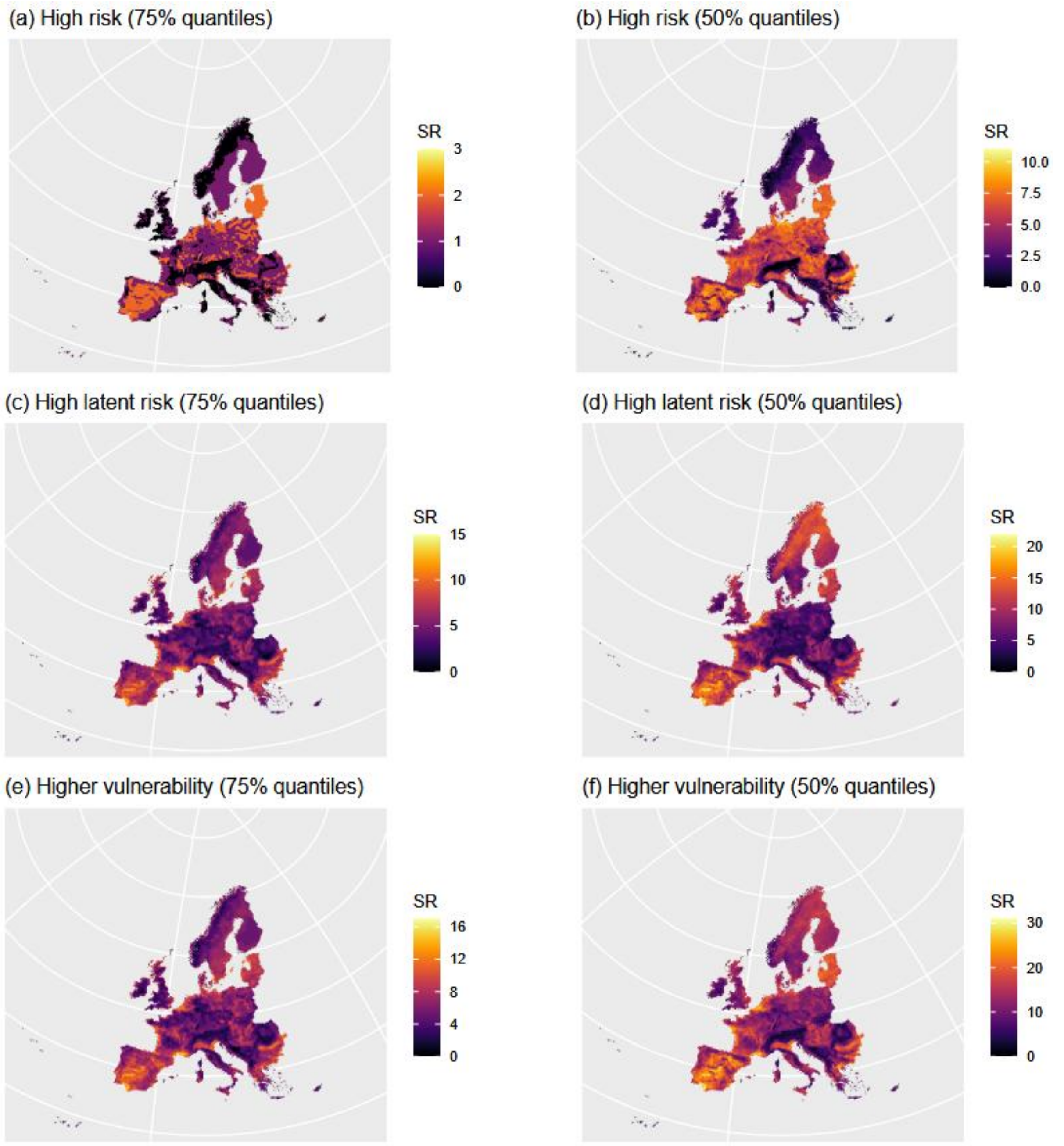


**Fig. S10.** Pseudo-species richness maps for species classified as high risk and high latent risk, and for the two threshold values used to classify species into risk categories (75% and 50% quantiles): **(a)** & **(b)** high risk; **(c)** & **(d)** high latent risk species; **(e)** & **(f)** when grouping together high risk and high latent risk (i.e., species with higher relative vulnerability).

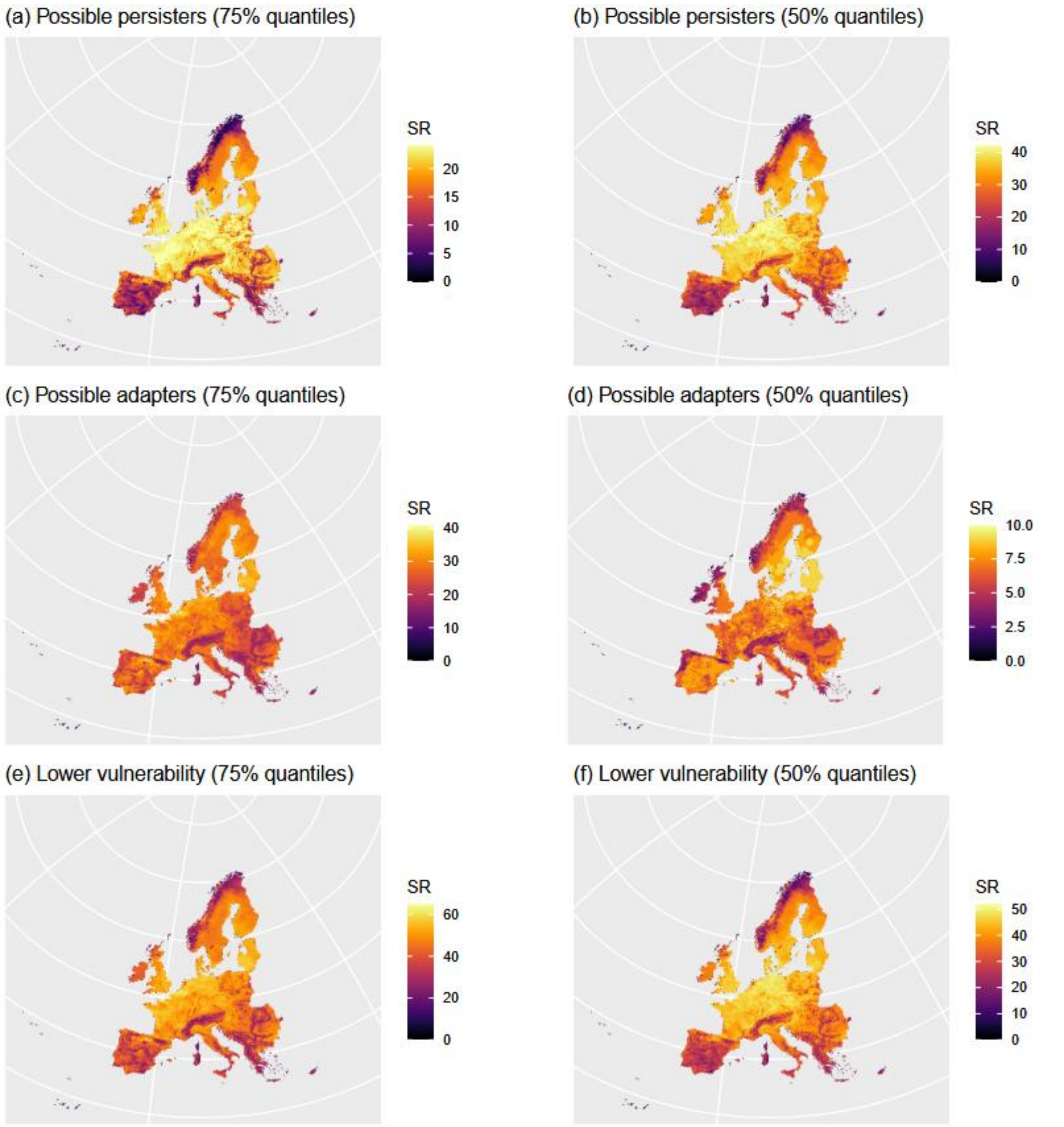


**Fig. S11.** Pseudo-species richness maps for species classified as possible persisters and possible adapters, and for the two threshold values used to classify species into risk categories (75% and 50% quantiles): **(a)** & **(b)** possible persisters; **(c)** & **(d)** possible adapters; **(e)** & **(f)** when grouping together possible adapters and possible persisters (i.e., species with lower relative vulnerability).

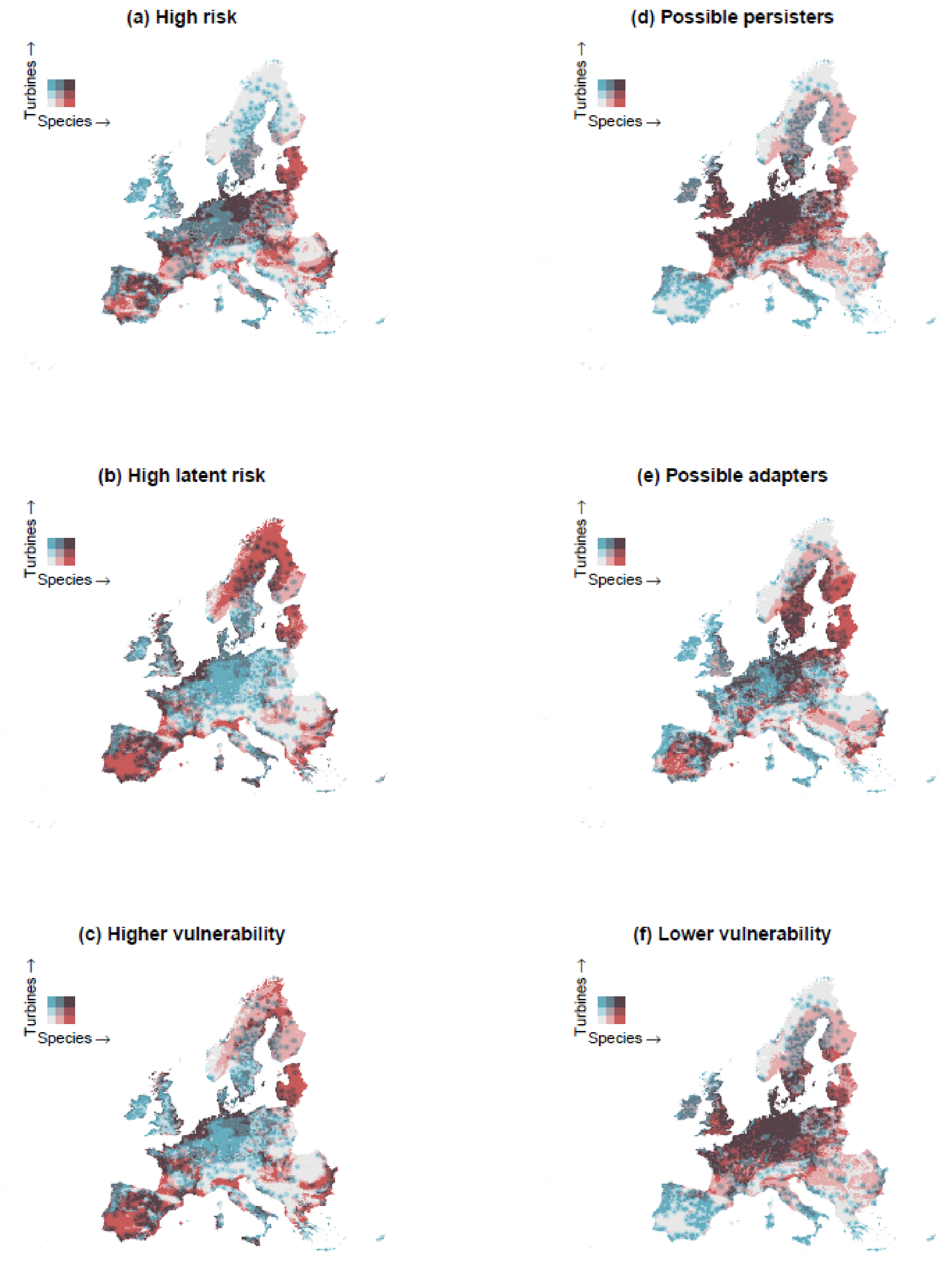


**Fig. S12.** Spatial overlap between suitable habitats for species in the different risk categories or groupings and current wind-turbine locations; **(a)** & **(b)** high risk and high latent risk species; **(c)** higher relative vulnerability (i.e., high risk and high latent risk species considered together); **(d)** & **(e)** possible persisters and possible adapters; **(f)** lower relative vulnerability (i.e., possible persisters and possible adapters considered together). Red areas harbour a larger number of species in a given risk category; blue areas harbour a larger number of turbines; purple areas indicate overlaps between areas of higher pseudo-species richness and higher turbine density. Here, risk categories were derived using the 50% quantiles of the distribution of the vulnerability and impact indices as thresholds.

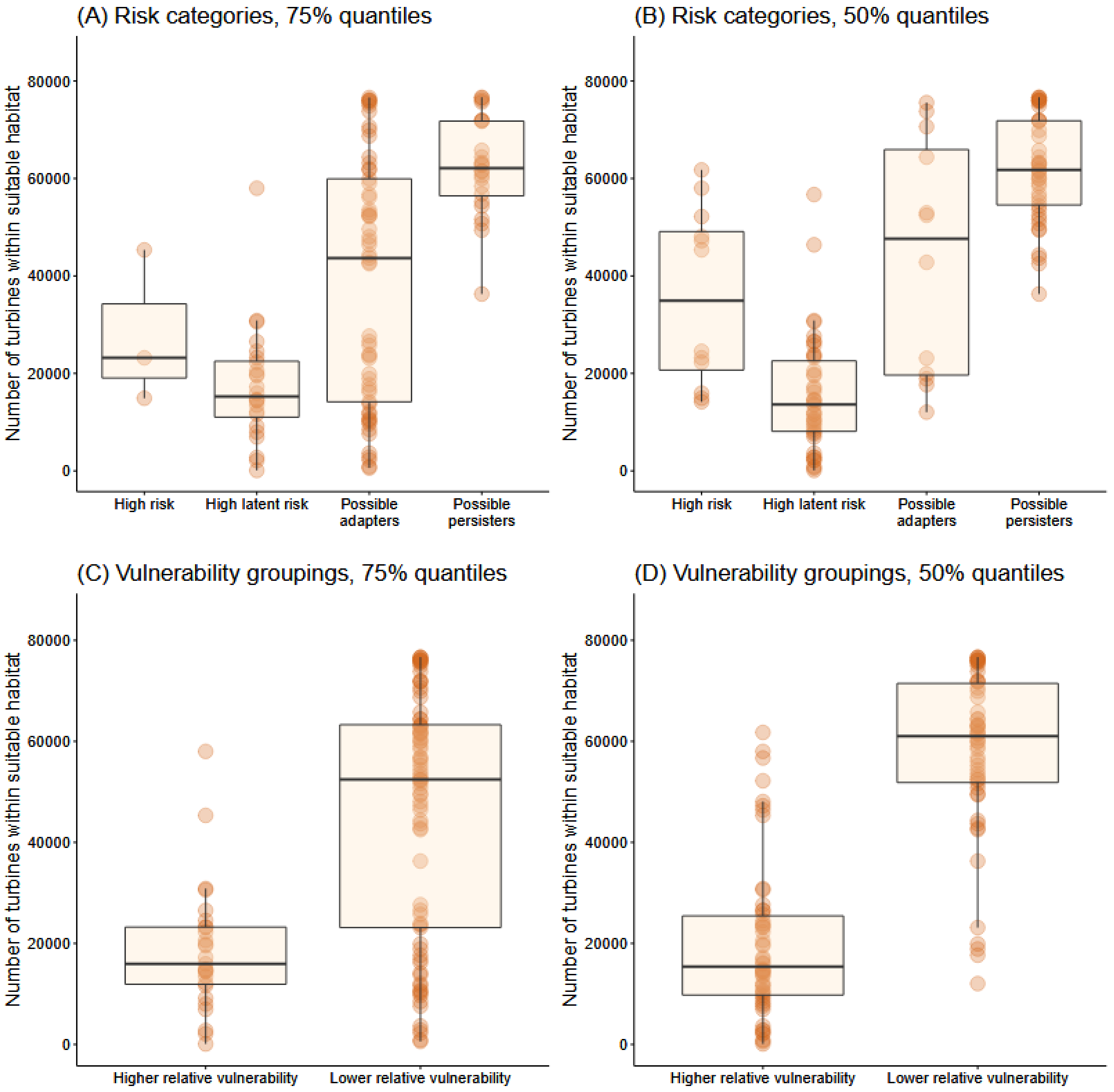


**Fig. S13.** Distribution of the number of turbines intersecting with the suitable areas of species in the different risk categories (**(A)** & **(B)**) and vulnerability groupings (**(C)** & **(D)**). Each dot represents a species.

#### S10. Assessment of potential biases in the fitted data

We conducted two additional analyses to examine the evidence for potential biases in the fitted data which could affect reported fatality counts and therefore estimated collision-mortality rates. Such biases could arise from (1) differences in search protocols among the included studies, or (2) differences in detectability among species.

(1) We investigated whether aspects of the experimental design that reflected survey effort in each study had an effect on collision-mortality rates. The variables pertaining to survey effort in each study, that were most widely available from the data, and that could influence the reported number of fatalities, were buffer area and number of monitoring days. Buffer area referred to the search area covered in each study, while the number of monitoring days described the total number of days monitored each year the study was conducted (note that this was different from the overall study duration in years; Thaxter et al. (2017)).
We fitted a model investigating whether collision-mortality rates were associated with buffer area and with the number of monitoring days (this model was the same as the 'taxonomic model' with these two variables added as fixed effects). This model was fitted across 73 sources (from which buffer area and monitoring days were available) sampling 323 species in 63 locations, totaling 2,098 records. Across studies, buffer area and monitoring days were not significantly associated with collision-mortality rates (Fig. S14). There was, therefore, no evidence that these differences in experimental design among the studies included influenced the estimated collision-mortality rates.

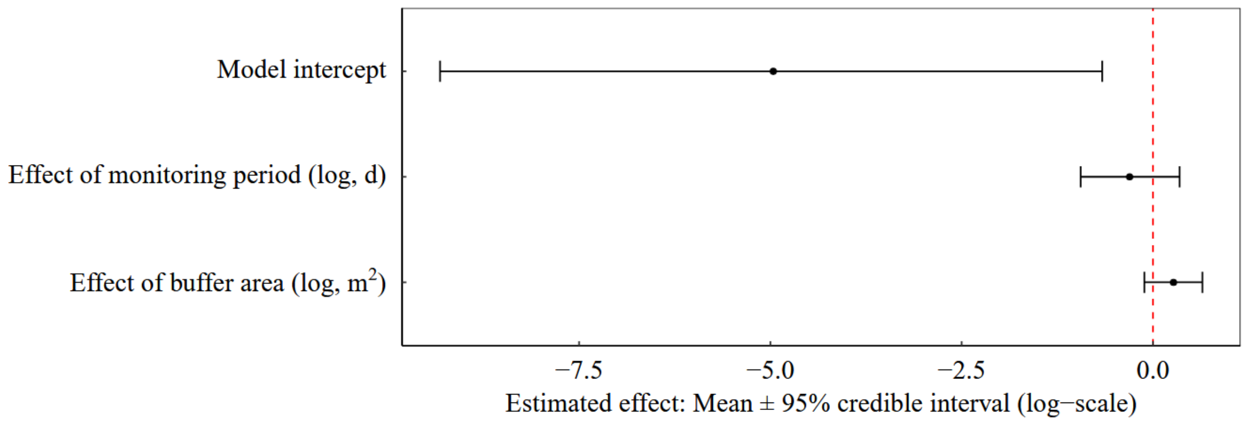


**Fig. S14.** Estimated effects of buffer area and of number of monitoring days on collision-mortality rates. We fitted a model similar to the 'taxonomic model' (see main text), with buffer area (log-transformed) and the number of monitoring days (log-transformed) added as fixed effects.

(2) We investigated whether body mass had an effect on collision-mortality rates. The rationale was that bigger species might be more detectable than smaller species, therefore biasing the reported number of fatalities towards bigger species (although all included studies were corrected for detectability in some way, see Main text, Results 3.1). We assessed whether there was evidence of such a detection bias in the fitted data by fitted a model similar to the 'trait model', whereby body mass (log-transformed) was included as the only species-level fixed effect. Body mass information were gathered from the AVONET database (Tobias et al.,

2022). This model was fitted on 328 species (for which body mass information was available) sampled across 81 sources and 69 locations, totaling 2,182 records. The effect of body mass was not significant (estimated effect: -0.06 [-0.12; 0.01]; Fig. S15), showing that on average, bigger species were not more likely to have higher collision mortality rates than smaller species.

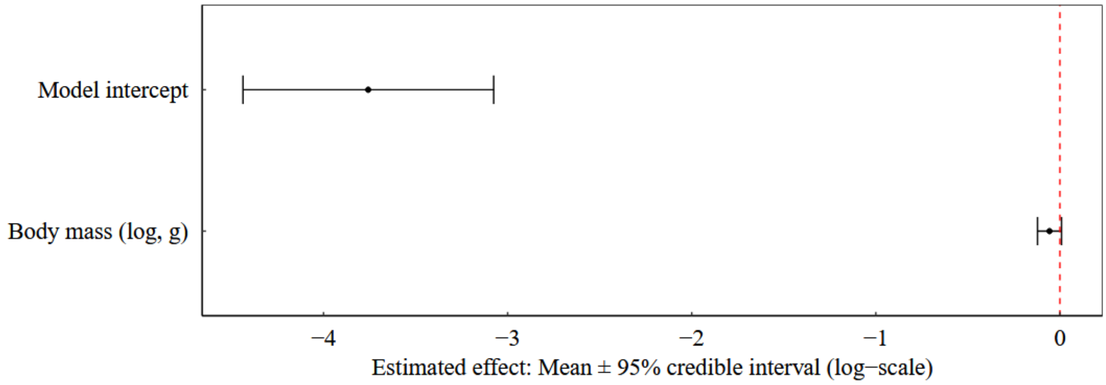


**Fig. S15.** Estimated effects of body mass on collision-mortality rates. We fitted a model similar to the ‘trait model’ (see main text), with body mass (log-transformed) included as the only species-level fixed effect fixed effects.

#### Supplementary references


Foden, W. B., Young, B. E., Akçakaya, H. R., Garcia, R. A., Hoffmann, A. A., Stein, B. A., Thomas, C. D., Wheatley, C. J., Bickford, D., Carr, J. A., Hole, D. G., Martin, T. G., Pacifici, M., Pearce-Higgins, J. W., Platts, P. J., Visconti, P., Watson, J. E. M., & Huntley, B. (2019). Climate change vulnerability assessment of species. *WIREs Climate Change*, *10*(1), e551. https://doi.org/10.1002/wcc.551

Massicotte, P., South, A., & Hufkens, K. (2023). *rnaturalearth: World Map Data from Natural Earth* (Version 1.0.1) [Computer software]. https://cran.r-project.org/web/packages/rnaturalearth/index.html

Santangeli, A., Butchart, S. H. M., Pogson, M., Hastings, A., Smith, P., Girardello, M., & Moilanen, A. (2018). Mapping the global potential exposure of soaring birds to terrestrial wind energy expansion. *Ornis Fennica*, *95*(1), Article 1. https://doi.org/10.51812/of.133925

Shiomi, K. (2022). Possible link between brain size and flight mode in birds: Does soaring ease the energetic limitation of the brain? *Evolution*, *76*(3), 649–657. https://doi.org/10.1111/evo.14425

Thaxter, C. B., Buchanan, G. M., Carr, J., Butchart, S. H. M., Newbold, T., Green, R. E., Tobias, J. A., Foden, W. B., O'Brien, S., & Pearce-Higgins, J. W. (2017). Bird and bat species' global vulnerability to collision mortality at wind farms revealed through a trait-based assessment. *Proceedings of the Royal Society B: Biological Sciences*, *284*(1862), 20170829. https://doi.org/10.1098/rspb.2017.0829

Tobias, J. A., Sheard, C., Pigot, A. L., Devenish, A. J. M., Yang, J., Sayol, F., Neate-Clegg, M. H. C., Alioravainen, N., Weeks, T. L., Barber, R. A., Walkden, P. A., MacGregor, H. E. A., Jones, S. E. I., Vincent, C., Phillips, A. G., Marples, N. M., Montaño-Centellas, F. A., Leandro-Silva, V., Claramunt, S., … Schleuning, M. (2022). AVONET: Morphological, ecological and geographical data for all birds. *Ecology Letters*, *25*(3), 581–597. https://doi.org/10.1111/ele.13898

Vehtari, A., Gelman, A., & Gabry, J. (2017). Practical Bayesian model evaluation using leave-one-out cross-validation and WAIC. *Statistics and Computing*, *27*(5), 1413–1432. https://doi.org/10.1007/s11222-016-9696-4

Watanabe, Y. Y. (2016). Flight mode affects allometry of migration range in birds. *Ecology Letters*, *19*(8), 907–914. https://doi.org/10.1111/ele.12627
